# Ribosome-associated quality control of aberrant protein production during amino acid limitation

**DOI:** 10.64898/2026.01.14.699605

**Authors:** Alicia M. Darnell, Christopher Chidley, Victoria Paradise, Danica S. Cui, Kristian Davidsen, Sarah C. Lincoln, Keene L. Abbott, Ryan Elbashir, Campbell P. Vander Heiden, Peter K. Sorger, Lucas B. Sullivan, Joseph H. Davis, Matthew G. Vander Heiden

## Abstract

Amino acids can become limiting for protein synthesis through depletion of charged tRNAs, leading to ribosome stalling and disruption of translation elongation at specific codons. To assess whether this is a mechanism by which amino acid availability can directly influence gene expression, we designed a reporter library to measure translation disruption across all sense codons in the context of amino acid limitations. We found that arginine limitation consistently impairs translation at the arginine codon AGA, resulting in synthesis of proteins from endogenous transcripts. In contrast, GCN2 pathway activation suppresses translation disruption following depletion of most other amino acids. Genome-wide screens revealed that the ribosome quality control trigger (RQC-T) and RQC pathways, which resolve ribosome collisions on defective mRNAs, catalyze ribosome splitting and premature fall-off in response to arginine depletion. Additionally, the E3 ubiquitin ligase RNF14, recently shown to clear ribosome A-site obstructions, promotes translation disruption through both ribosome fall-off and frameshifting during arginine limitation. Together, these data show that the RQC machinery is engaged by tRNA-limited ribosomes and identify a new role for RNF14 as a regulator of translation upon arginine limitation.

## INTRODUCTION

Amino acid availability can directly integrate cellular metabolism with gene expression when it limits protein synthesis. Amino acid depletion can affect the activity of nutrient-sensing kinases such as mTOR and GCN2 that regulate rates of ribosome initiation^1^; however, amino acid depletion can also cause ribosome stalling through the depletion of charged tRNAs^2–6^. Ribosome stalling has been proposed to act as both an ancient mechanism of gene regulation at specific synonymous codons, and a source of translational stress. It has been linked to biofilm formation^7^, neurodegeneration^8^, cancer metastasis^9^, tumor progression^5^, and mitochondrial function^10^. However, the environmental conditions that trigger ribosome stalling, and the codons at which it occurs, remain incompletely understood. Further, how ribosome stalling is resolved and regulated, and whether it results in proteome alterations, is an area of active research.

Protein synthesis rate is generally limited by initiation rate^11–13^, but several mechanisms have been proposed by which ribosome stalling can directly alter protein production by affecting elongation. For example, collisions between the stalled and trailing ribosomes can cause translation elongation disruption^14,15^. On damaged or defective mRNAs, collisions trigger ribosome ubiquitination^16–19^ and subsequent ribosome splitting through the ribosome quality control trigger complex (RQC-T)^20,21^. This is followed by downstream nascent polypeptide degradation via the ribosome quality control (RQC) pathway^22–24^ and can be coupled to mRNA degradation by the no-go decay (NGD) pathway^25,26^. However, whether these pathways act on ribosomes that are reversibly slowed on intact mRNAs by reduced charged tRNA abundance is less clear. Stable nascent polypeptide products have been detected downstream of translation disruption in leucine deprived cells^2^. Further, specific amino acid or charged tRNA limitations can also lead to ribosome frameshifting^4,27–29^ and amino acid misincorporation^30–33^ to alter protein output.

As a systematic approach to determine whether depletion of each amino acid leads to ribosome stalling and translation elongation disruption, and to probe the mechanisms and regulators involved, we designed a library of fluorescent reporters that measure translation disruption leading to aberrant protein production at each sense codon. Depletion of several amino acids, most notably arginine, triggered codon-specific ribosome stalling sufficient to cause translation disruption. In contrast, the integrated stress response through GCN2 kinase prevented translation disruption following depletion of most other amino acids. Arginine limitation disrupted ribosome elongation with high specificity at the arginine codon AGA, leading to reduced endogenous protein production that scaled with a gene’s AGA content. Genome-wide CRISPRi/a screens for regulators of ribosome stalling at AGA codons revealed that the RQC-T and RQC pathways catalyze ribosome splitting and nascent peptide quality control during arginine limitation. These screens also showed that arginine-limited translation disruption is regulated by the E3 ubiquitin ligase RNF14, recently implicated in the clearance of trapped ribosomal A-site proteins and RNA-protein crosslinks^34–37^. Taken together, this work highlights a role for RNF14 and the RQC pathway in shaping protein production in response to arginine limitation.

## RESULTS

### A survey of the effect of amino acid depletion at each codon reveals arginine AGA codons as a unique trigger for translation disruption

We designed a library of reporters to assess which amino acid limitation conditions and codons can trigger ribosome stalling and translation disruption. Flag-tagged YFP (Flag-YFP) was fused to a trimethoprim (TMP)-stabilizable E. coli dihydrofolate reductase-based degron domain (DHFR)^38,39^ using a linker containing query codons (Fig. 1A). Thus, limitation for an amino acid leading to translation disruption at the linker will generate a fluorescent Flag-YFP polypeptide lacking the degron domain, which can be detected by flow cytometry or western blotting^2^. To minimize translation disruption within YFP, we determined the codon that is most robustly translated for each amino acid either from published ribosome profiling data^2,5^, or by treating cells expressing each YFP codon variant with TMP and monitoring protein synthesis rate via fluorescence accumulation upon depletion of the cognate amino acid^2,39^ (File S1; Table 1). For amino acids encoded by only two codons, we tested all possible combinations of YFP and linker codon variants.

**Figure 1:**
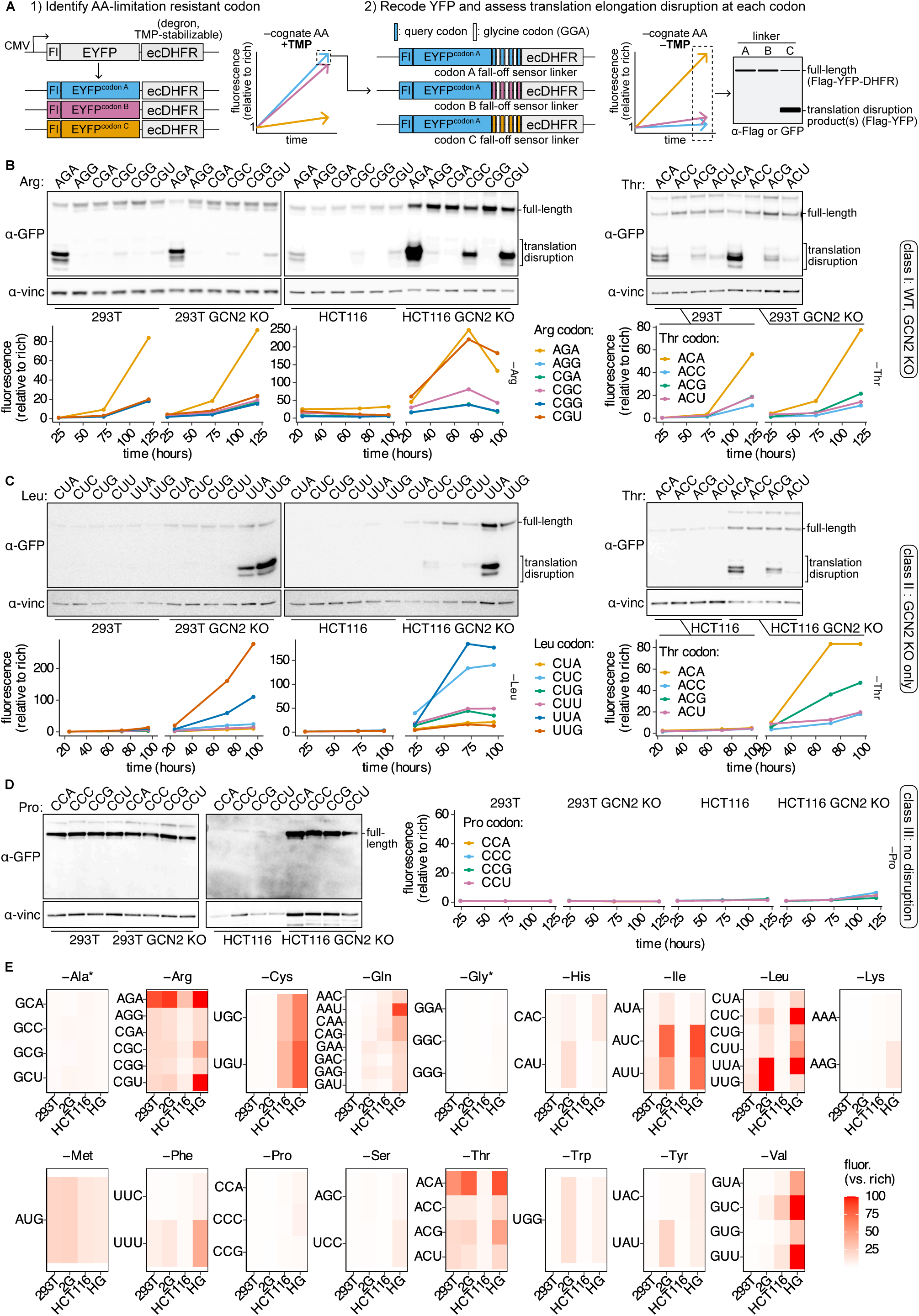
A survey of translation disruption across codons and conditions reveals that arginine AGA codons are a unique trigger. (A) Outline of translation disruption reporter survey across all codons. First, recoding the region upstream of the degron (ecDHFR^116,124^ or “DHFR”) with all possible codons for the amino acid (AA) of interest and adding TMP to prevent degradation and measure YFP synthesis rate^116^ upon limitation for that AA was used to determine the most AA limitation-resistant codon. Next, engineering linkers downstream of recoded AA limitation-resistant-codon YFP (Flag-YFP-4xNNN-DHFR) allows detection of translation disruption via YFP accumulation in the absence of TMP by either flow cytometry or western blotting. Alternatively, for AAs with only 2 codons, all 4 combinations of YFP/linker codon variants were tested to identify the combination most able to assess translation disruption. (B-D) Western blot to detect translation disruption product accumulation during limitation for examples of class I (arginine, threonine in 293T cells), class II (leucine, threonine in HCT116 cells), or class III (proline) amino acids, and flow cytometry to detect reporter fluorescence change (Flag-YFP-4x[codon]-DHFR) upon the same conditions in wildtype (WT) and GCN2 knockout (KO) 293T or HCT116 cells as indicated. For blots, arginine was limited for 5 days in 293T and 3 days in HCT116 cells. Threonine was limited for 5 days in 293T cells and 4 days in HCT116 cells. Leucine was limited for 4 days in both cell lines. Proline was limited for 5 days in both cell lines. Full-length and translation disruption reporter products are indicated (α-GFP antibody used to detect YFP, α-vinc = α-vinculin as a loading control). (E) Heatmaps depicting reporter fluorescence change upon limitation for the indicated amino acid at the indicated codon across WT and GCN2 KO 293T (293T vs “2G”) and HCT116 (HCT116 vs “HG”) cells as indicated. Color scale is capped at a maximum value of 100. Notes on amino acid limitation conditions: −Ala*: mitochondrial pyruvate carrier (MPC) inhibitor UK5099 in −glucose used to deplete alanine; −Gly*: −Serine,−glycine used to deplete glycine. −Glutamine: used to deplete asparagine, aspartate, glutamine, and glutamate.

**TABLE 1:** Translation disruption reporter library design for survey.

Ribonucleotide abbreviations: N = A,C,G,U ; R = A, G (purines) ; Y = C, U (pyrimidines). AA = amino acids; KO = knockout; KD = knockdown; MPCi = UK5099, mitochondrial pyruvate carrier inhibitor.
| **Amino Acid** | **Limitation Condition** | **YFP variants tested?** | **YFP variant used** | **Linkers tested ?** | **"In between" codons in linker** | **Rationale** |
| --- | --- | --- | --- | --- | --- | --- |
| Alanine | Glucose + MPCi | all | GCU | all | GGA | Pilot test comparing YFP synthesis rates |
| Cysteine | Cysteine +- ferrostatin | wild-type Flag-EYFP: 2 UGC | UGC | all | GGA | Low representation in YFP |
| Aspartate | Glutamine | all | GAC | all | GGA | Tested all possible combinations |
| Glutamate | Glutamine | wild-type Flag-EYFP: 15 GAG, 1 GAA | wild-type YFP: 15xGAG, 1xGAA | all | GGA | Prior knowledge ribosome profiling (VanInsberghe et al 2021) |
| Phenylalanine | Phenylalanine | all | UUC | all | GGA | Tested all possible combinations |
| Glycine | Serine, glycine | all | GGC | all | GAC | Pilot test comparing YFP synthesis rates |
| Histidine | Histidine | all | CAC | all | GGA | Tested all possible combinations |
| Isoleucine | Isoleucine | all | AUA | all | GGA | Pilot test comparing YFP synthesis rates |
| Lysine | Lysine | all | AAA | all | GGA | Tested all possible combinations |
| Leucine | Leucine | CUA | CUA | all | GGA | Prior knowledge ribosome profiling + pilot test comparing YFP synthesis rates (Darnell et al 2018) |
| Methionine | Methionine | wild-type Flag-EYFP: 6 AUG | AUG | all | GGA | Only 1 codon |
| Asparagine | Glutamine | all | AAC | all | GGA | Tested all possible combinations |
| Proline | Proline +- arginine or PYCR triple KO | all | CCG | all | GGA | Pilot test comparing YFP synthesis rates |
| Glutamine | Glutamine, all AA | all | CAA | all | GGA | Tested all possible combinations |
| Arginine | Arginine | CGG | CGG | all | GGA | Prior knowledge ribosome profiling + pilot test comparing YFP synthesis rates (Darnell et al 2018) |
| Serine | Serine +/- glycine, rotenone, or PHGDH KD | AGC | AGC | AGC, UCC | GAC | Prior knowledge ribosome profiling + pilot test comparing YFP synthesis rates (Banh et al 2020) |
| Threonine | Threonine | all | ACC | all | GGA | Pilot test comparing YFP synthesis rates |
| Valine | Valine | all | GUG | all | GGA | Pilot test comparing YFP synthesis rates |
| Tryptophan | Tryptophan | wild-type Flag-EYFP: 1 UGG | UGG | all | GGA | Low representation in YFP |
| Tyrosine | Tyrosine | all | UAC | all | GGA | Tested all possible combinations |

The reporter library was expressed in HEK293T (“293T”) and HCT116 colorectal cancer cells, as these cells were previously demonstrated in ribosome profiling studies to exhibit ribosome stalling during limitation for some amino acids^2,40,41^. As the adaptive signaling response through GCN2 kinase suppresses translation initiation and thereby mitigates ribosome stalling and collisions^2,42^, we also expressed the reporter library in 293T and HCT116 GCN2 knockout (KO) cells (Fig. S1A) to assess the role of GCN2 in preventing translation disruption. We completely deprived cells for each amino acid to maximize the likelihood of ribosome stalling; these conditions generally reduced or arrested cell proliferation, particularly in GCN2 KO cells (Fig. S1B). For the nonessential amino acids proline, alanine, glycine, asparagine, aspartate, and glutamate that are not required for proliferation in culture we depleted metabolic precursors or generated auxotrophic cell lines using gene knockout to achieve intracellular depletion of those amino acids (Table 1).

For each reporter, we measured Flag-YFP accumulation by flow cytometry (Fig. 1B-E, Files S2,3) and confirmed the production of truncated polypeptides by western blotting (Fig. 1B-D, S1C, Files S4,5). Translation disruption was condition-specific, and three general patterns emerged; amino acids for which limitation caused translation disruption in at least one of the “wild-type” (WT) cells tested (termed “class I”; Fig. 1B), only in GCN2 KO cells (“class II”; Fig. 1C), or in neither genotype (“class III”; Fig. 1D; Table 2). Translation disruption was also synonymous codon-specific, consistent with prior observations in both *E. coli*^3,7,43,44^ and human cells^2,5^ (Fig. 1B-E; Table 2).

**TABLE 2:**
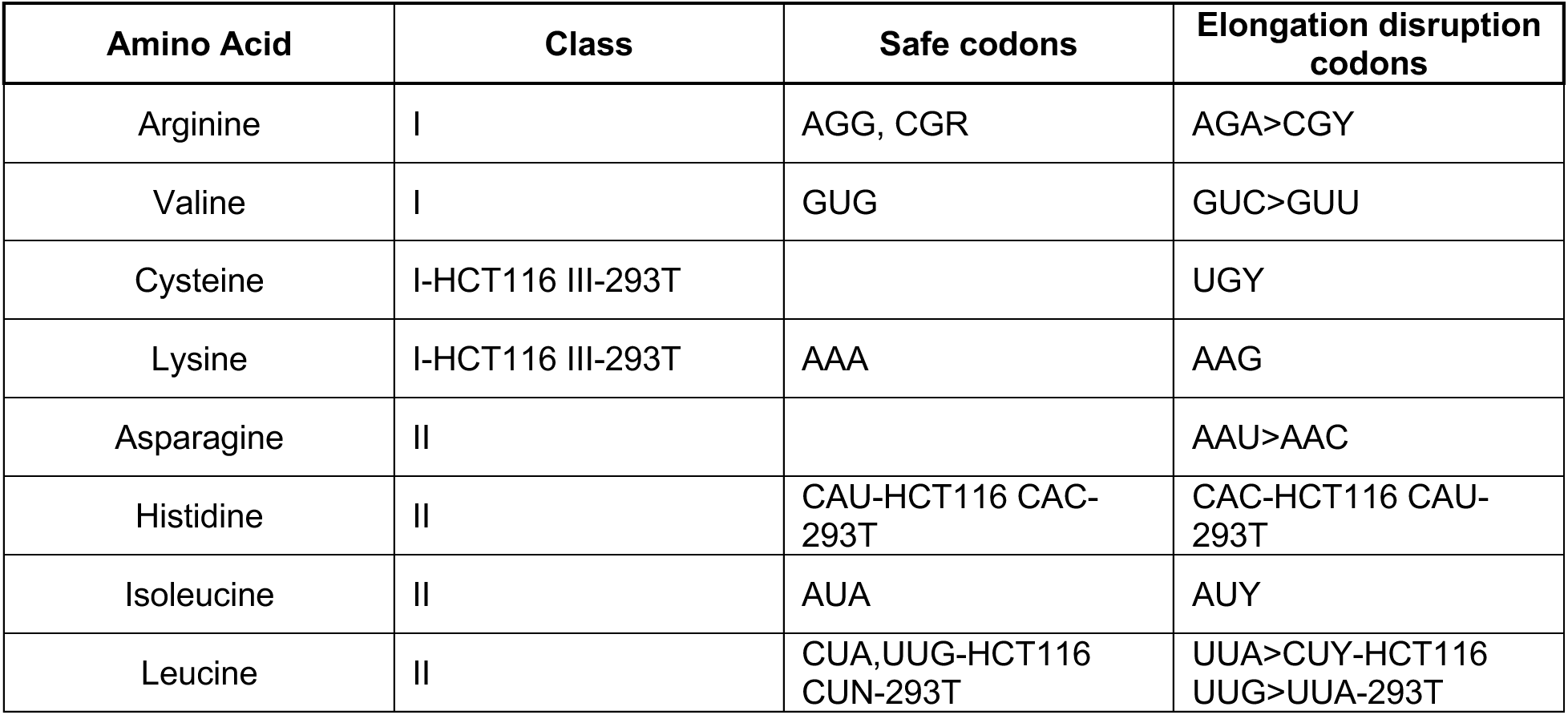
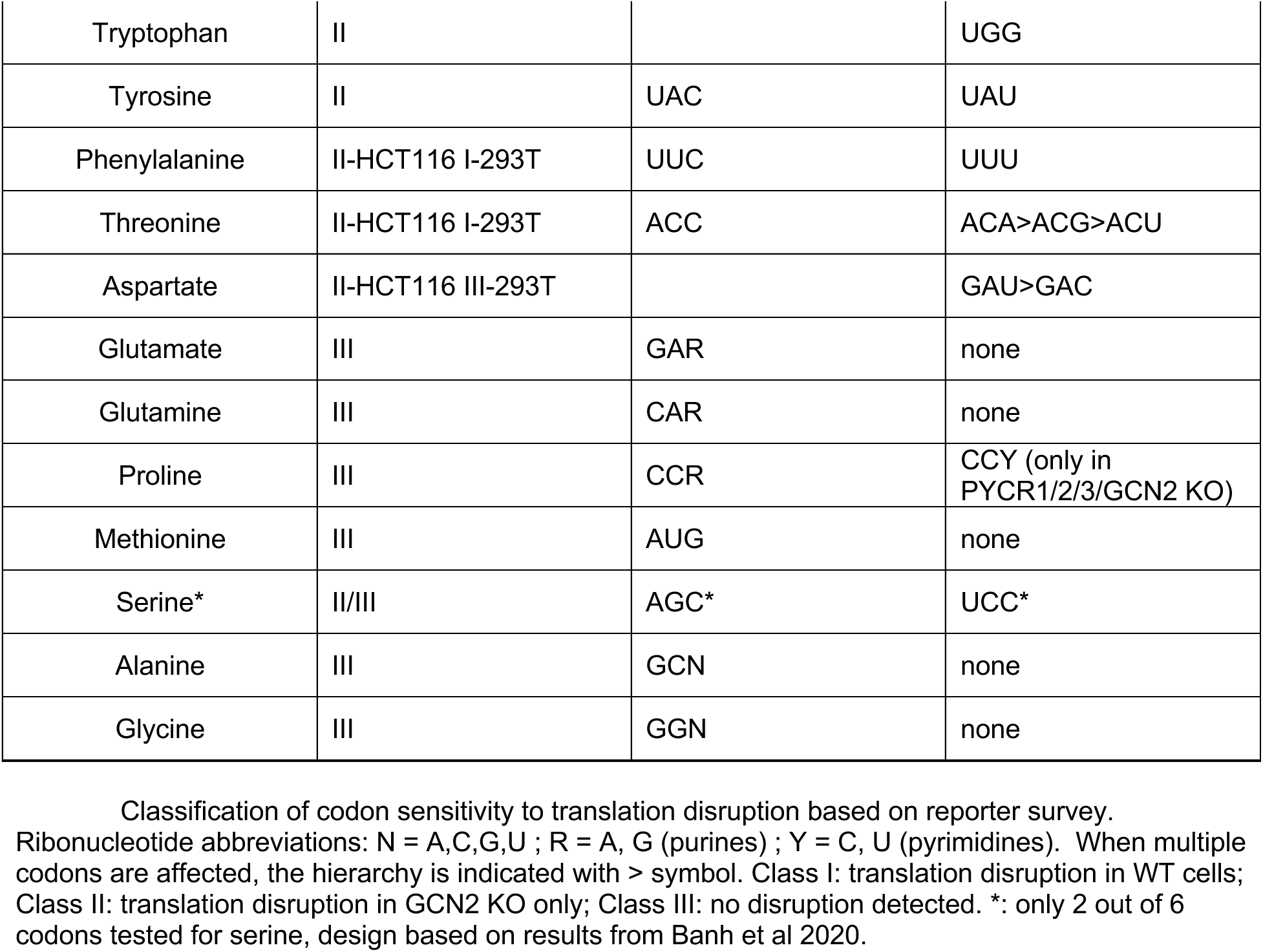
Classification of codon sensitivity to translation disruption.

based on reporter survey.
| Amino Acid | Class | Safe codons | Elongation disruption codons |
| --- | --- | --- | --- |
| Arginine | I | AGG, CGR | AGA>CGY |
| Valine | I | GUG | GUC>GUU |
| Cysteine | I-HCT116 III-293T |  | UGY |
| Lysine | I-HCT116 III-293T | AAA | AAG |
| Asparagine | II |  | AAU>AAC |
| Histidine | II | CAU-HCT116 CAC-293T | CAC-HCT116 CAU-293T |
| Isoleucine | II | AUA | AUY |
| Leucine | II | CUA,UUG-HCT116 CUN-293T | UUA>CUY-HCT116 UUG>UUA-293T |
| Tryptophan | II |  | UGG |
| Tyrosine | II | UAC | UAU |
| Phenylalanine | II-HCT116 I-293T | UUC | UUU |
| Threonine | II-HCT116 I-293T | ACC | ACA>ACG>ACU |
| Aspartate | II-HCT116 III-293T |  | GAU>GAC |
| Glutamate | III | GAR | none |
| Glutamine | III | CAR | none |
| Proline | III | CCR | CCY (only in PYCR1/2/3/GCN2 KO) |
| Methionine | III | AUG | none |
| Serine* | II/III | AGC* | UCC* |
| Alanine | III | GCN | none |
| Glycine | III | GGN | none |

Among all tested conditions, arginine limitation was the most consistent trigger of translation disruption in WT cells, though limitation for threonine, cysteine, valine, phenylalanine, and lysine led to detectable codon-specific translation disruption in at least one cell line (Fig. 1B,E; S1C, Files S2-5). Of the six arginine codons, AGA was the most sensitive, though translation disruption was also detectable at CGC and CGU codons, consistent with prior observations that acute arginine limitation can cause ribosome stalling at these codons^2,40,41^. To exclude translation disruption arising from poly-arginine tracts, as has been reported in yeast^45–47^, we tested reporters containing only 2 arginine codons separated by a flexible serine/glycine linker and still observed strong AGA-specific translation disruption across a panel of human cells (Fig. S1D). Translation disruption occurred at extracellular arginine levels at or below 2 μM (Fig. S1E). It did not depend on the “safe” arginine codon used to encode YFP or the degron, the N-terminal tag, the addition of TMP to stabilize any full-length reporter synthesized (Fig. S1F-G), or linker sequence context (Fig. S1H), indicating that the observed effect reflects codon-specific elongation failure rather than reporter design.

The GCN2 response prevented translation disruption during limitation for many amino acids. Limitation for leucine, tryptophan, isoleucine, histidine, and tyrosine caused minimal translation disruption in WT cells but led to robust codon-specific disruption in GCN2 KO cells (Fig. 1C,E; Files S2-5). Glutamine depletion also caused robust codon-specific translation disruption at the asparagine codon AAU in both GCN2 KO cell lines (Fig. S1I, Files S4,5) despite reducing intracellular levels of glutamine, glutamate, and aspartate as well as asparagine (Fig. S1J), consistent with prior reports that intracellular asparagine limits proliferation upon glutamine restriction^48,49^.

Several amino acid limitation conditions caused little or no translation disruption. Proline codons had previously been found to display mild ribosome stalling in clear cell renal carcinoma tumors^50^, but we detected minimal translation disruption in 293T or HCT116 cells in the absence of extracellular proline (Fig. 1D,E; Files S2-5). Limitation for arginine and glutamine, substrates for proline synthesis, in addition to proline, also caused minimal translation disruption (Fig. S1K). To confirm this was due to proline synthesis by these cells, we knocked out all three PYCR isoforms (PYCR TKO) in WT and GCN2 KO 293T cells to generate proline auxotrophs (Fig. S1L,M). GCN2 KO PYCR TKO, but not WT PYCR TKO cells displayed translation disruption at CCC and CCU proline codons upon proline limitation (Fig. S1N).

Likewise, serine limitation did not lead to translation disruption in 293T or HCT116 cells (Fig. 1E, Files S2-5), likely due to sufficient serine synthesis. To confirm this, we used the respiration inhibitor rotenone to restrict serine synthesis^51^; this led to translation disruption at UCC codons in GCN2 KO but not WT HCT116 cells during serine limitation (Fig. S1O), in line with ribosome stalling previously observed at UCC/U codons in pancreatic ductal adenocarcinoma cells with low expression of the serine synthesis enzyme PHGDH^5^. In MDA-MB-231 breast cancer cells, which express low levels of PHGDH^52^, translation disruption was also not observed when GCN2 was intact (Fig. S1P). Interestingly, despite intact GCN2, translation disruption occurred in the PHGDH-expressing primary mouse breast cancer cell line (MS-1579) following PHGDH silencing by CRISPRi^52^ (Fig. S1Q), suggesting that cells with low PHGDH may rely more on the GCN2 response to prevent translation disruption upon serine deprivation. Translation disruption did not occur at glycine codons upon limitation for serine and glycine, or alanine codons upon limitation for glucose with inhibition of the mitochondrial pyruvate carrier (MPC), a condition that leads to alanine auxotrophy^53–55^(Fig. 1E).

Finally, cysteine limitation caused translation disruption in HCT116 but not 293T cells (Fig. 1E; Fig. S1R). Cell-type specific metabolic or signaling responses may explain this difference. During cysteine limitation, 293T cells underwent ferroptosis (Fig. S1S), while HCT116 cells are relatively ferroptosis resistant^56^; divergent translation disruption responses may depend on whether levels of Cys-tRNA^Cys^ or the antioxidant glutathione^57^ are affected.

While translation disruption was codon-specific, we noticed that full-length fluorescent YFP-DHFR “background” signal accumulated upon limitation for many amino acids (Fig. 1B-E, Files S1-5). This was not due to impaired proteasome activity, as global protein turnover was unaffected by arginine limitation (Fig. S1T). Instead, we observed that mRNA levels of reporter transgenes increased during amino acid deprivation (Fig. S1U), and did so proportional to the severity of translation disruption observed (Fig. S1V–Z), consistent with previous reports that lentivirally integrated transgenes are transcriptionally induced under conditions of translational stress^58^. Importantly, this effect did not account for codon-specific translation disruption, which was directly confirmed by the accumulation of truncated polypeptides detected by western blotting.

Overall, these results highlight that ribosome stalling severe enough to cause translation disruption in response to amino acid limitation depends on the specific amino acid, synonymous codon, and cell type, and is often prevented by the adaptive GCN2 response. In the cell lines tested here, arginine limitation stood out as the most consistent metabolic trigger of ribosome stalling and translation disruption at AGA codons.

### Isoacceptor-specific charged tRNA depletion underlies codon-specific translation disruption

Amino acid limitation has been proposed to cause uneven depletion of charged isoacceptor tRNAs in bacteria^3,7,43,44^ and in human cells, leading to codon-specific ribosome stalling^2^. To test whether this accounts for codon-specific translation disruption in the cells surveyed, we performed charged tRNA sequencing (Fig. 2A)^59,60^ in both WT and GCN2 KO 293T and HCT116 cells during limitation for 11 amino acids representing all three classes identified in the reporter survey. We observed progressive depletion of cognate charged tRNA levels during limitation for all amino acids tested except serine, glycine, and glutamine (Fig. S2A), consistent with intracellular synthesis being sufficient to prevent translation disruption in these cases (Fig. 1E; S1I,O,Q). GCN2 KO exacerbated tRNA charging loss in response to some, but not all amino acid depletion conditions (Fig. S2A).

**Figure 2:**
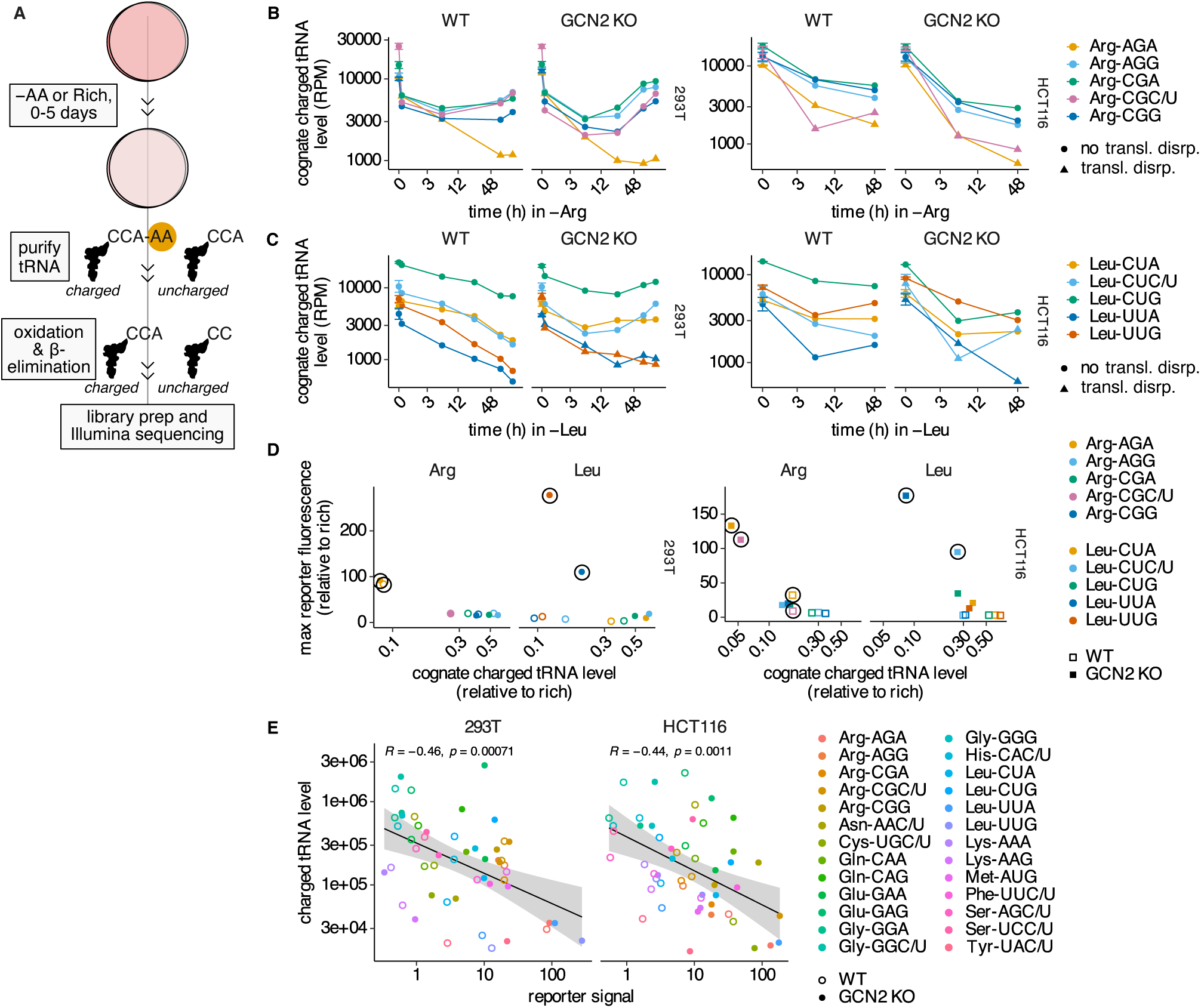
Isoacceptor-specific charged tRNA depletion underlies codon-specific translation disruption. (A) Schematic depicting charged tRNA sequencing method. (B-C) Charged tRNA levels (reads per million; RPM) for the arginine (B) and leucine (C) tRNAs cognate to the indicated codons upon limitation for the corresponding amino acid in wild-type (WT) and GCN2 KO 293T or HCT116 cells as indicated. Error bars are only present at t=0 and reflect n=3 biological replicate measurements. Codons at which translation disruption (transl. disrp.) was detected by western blotting using the reporter described in Fig. 1 are indicated with a triangle. (D) Maximum reporter fluorescence signal measured at each arginine/leucine codon versus the cognate charged tRNA level at the last time point measured for WT and GCN2 KO 293T or HCT116 cells as indicated. Codons at which translation disruption was detected by western blotting are indicated with a surrounding black circle. (E) Correlation between reporter fluorescence signal and cognate charged tRNA level at the same time point measured across WT and GCN2 KO 293T or HCT116 cells (log_10_ scaled axes). Spearman correlation coefficient (R) and associated p-value are shown on the plot. (B-E) For tRNAs that decode more than one codon through wobble base pairing, multiple codons are indicated in the legend.

The codon-specific pattern of translation disruption observed during limitation for an amino acid could be explained by differential reduction of specific charged tRNA levels in most cases. During arginine limitation, charged levels of Arg-tRNA^UCU^, which decodes AGA, were lower than other Arg-tRNAs, consistent with translation disruption at AGA codons (Fig. 2B, Fig. 1E). Depletion of charged Arg-tRNA^UCU^ was due to both lower fractional charging and reduced overall Arg-tRNA^UCU^ levels relative to other Arg-tRNAs upon arginine limitation (Fig. S2B,C), consistent with a recent report in colorectal cancer cells^61^. In HCT116 cells, charged Arg-tRNA^A/ICG^, which decodes CGC and CGU codons via inosine modification at the wobble position, was also depleted, in line with translation disruption observed at those codons (Fig. 2B, Fig. 1E). Our finding that translation disruption is less severe at CGC/U codons in 293T cells was explained by higher total tRNA^A/ICG^ levels despite its low percent charging (Fig. S2B,C).

Charged tRNA levels also explained the divergent impact of leucine limitation on leucine codons between 293T and HCT116 cells (Fig. 2C); both cell lines experienced translation disruption at UUA codons, but in 293Ts translation was disrupted at UUG codons, while in HCT116 cells, CUC/U codons were affected (Fig. 1C,E). In 293T cells, leucine limitation led to the largest decline in charged Leu-tRNA^CAA^ and Leu-tRNA^UAA^ (Fig. 2C), which decode the affected UUG and UUA codons, respectively. In HCT116 cells, loss of charged tRNA was correspondingly greatest for Leu-tRNA^AAG^, which decodes CUC/U codons via wobble base pairing, and Leu-tRNA^UAA^ (Fig. 2C). Total tRNA levels were lowest amongst the Leu-tRNAs for Leu-tRNA^UAA^ (Fig. S2D), perhaps explaining the sensitivity of UUA to translation disruption in both cell lines as suggested in the context of ribosome frameshifting^4^.

Integrating information about tRNA charging with tRNA modification state corroborated other conclusions from the reporter survey. In GCN2 KO but not WT cells, glutamine limitation led to sustained depletion of asparagine charged tRNA relative to rich conditions (Fig. S2F), consistent with GCN2 KO-specific translation disruption at asparagine codons (Fig. 1E, S1I; files S2-5). The asparagine codons AAC and AAU are both decoded by Asn-tRNA^GUU^, raising the question of why translation disruption was primarily evident at AAU codons. Decoding of NA<u>U</u> codons is enhanced by a guanine to queuosine (Q) anticodon modification^62,63^, which requires queuine uptake^64^, and queuine availability is likely reduced by the use of dialyzed serum^65^ for preparation of media lacking amino acids. Indeed, we observed less G^34^-modified Asn-tRNA^GUU^ upon glutamine limitation (Fig. S2G), and a higher likelihood of translation disruption at other <u>N</u>AU vs <u>N</u>AC wobble decoded pairs including tyrosine, histidine, and aspartate (Files S2-5).

Though absolute charged tRNA levels were a better predictor of translation disruption than percent charging, translation disruption generally occurred at tRNA charging levels less than ∼25% (Fig. 2D). GCN2 KO cells were more prone to translation disruption than WT cells even at similar charged tRNA levels, particularly at leucine codons (Fig. 2C,D); overactive ribosome initiation in these cells stimulates ribosome collisions^2,42^, which trigger translation disruption during other types of translational stress^15^. Importantly, lower charged tRNA levels were associated with an increased translation disruption reporter signal, although low levels of charged tRNA were not always sufficient to cause translation disruption (Fig. 2E).

### Translation disruption at AGA codons reduces endogenous protein production

We next sought to explore whether translation disruption at AGA codons in response to arginine limitation is sufficient to alter protein production. We first used a modified YFP-DHFR reporter in which all arginine codons were altered to each of the six arginine variants. YFP synthesis was impeded by CGC, CGU, and AGA codons after acute arginine limitation as previously observed^2^, but at longer times AGA codons emerged as the strongest disruptor of protein production (Fig. S3A).

To quantify the impact of translation disruption on endogenous protein production, we performed pulsed-SILAC (stable isotope labeling with amino acids in cell culture) proteomics to measure gene-specific protein synthesis rates in WT and GCN2 KO 293T cells limited for arginine or leucine (Fig. 3A). We reasoned that if translation disruption at AGA codons impaired elongation on endogenous transcripts, genes enriched for AGA codons would exhibit reduced synthesis rates during arginine limitation (Fig. S3B). Consistent with this hypothesis, AGA frequency, but not the frequency of any other arginine codon, was inversely correlated with the change in protein synthesis rate upon arginine limitation in WT and GCN2 KO cells (Fig. 3B,C, Fig. S3C). No such relationship was observed during leucine limitation.

**Figure 3:**
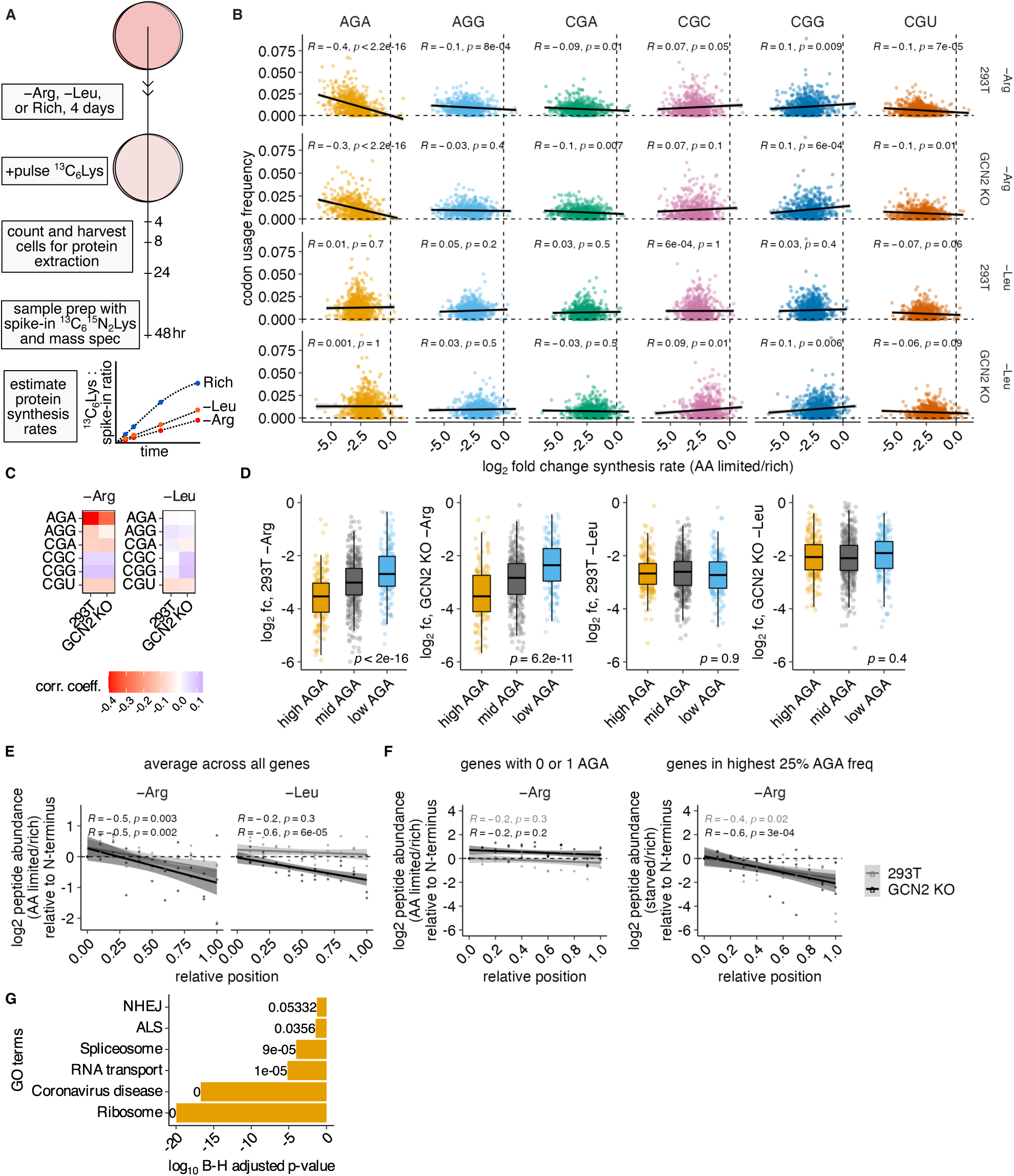
Translation disruption at AGA codons reduces endogenous protein production. (A) A schematic depicting the experimental approach for use of stable isotope labeling of amino acids in culture (SILAC) to measure protein synthesis rates upon arginine or leucine limitation. (B) Transcript codon usage frequency for each arginine codon plotted against the log_2_ fold-change in protein synthesis rate for the corresponding protein upon limitation for arginine or leucine for 4 days in wild-type and GCN2 KO 293T cells as indicated. Regression line, Spearman correlation coefficient (R) and associated p-value are shown on each plot. Note that 5 outlier points with log_2_ fold-change (fc) below −7.5 are not plotted. (C) A heatmap of associated Spearman correlation coefficients (corr. coeff.) from (B). (D) Boxplot showing the distribution of log_2_ fc in protein synthesis rate upon limitation for arginine or leucine for 4 days in wild-type and GCN2 KO 293T cells versus the transcripts binned by AGA usage frequency (high = top 25%, mid = 25-75%, low = bottom 25%). P-values represent one-way ANOVA tests comparing log₂ fold-change distributions across AGA classes (high, mid, low) within each condition. (E,F) Log_2_ fc in average peptide abundance upon limitation for arginine or leucine for 5 days including after ^13^C_6_Lys pulse for the last 24 hours, plotted against the binned fractional position of the peptide from 0 (N-terminus) to 1 (C-terminus), normalized to abundance at the N-terminus, for all peptides (E) or peptides from transcripts with ≤1 AGA codons or in the top 25% of AGA usage frequency (F) in wild-type and GCN2 KO 293T cells as indicated. (G) GO term enrichment analysis for the top 118 candidate AGA-regulated genes (NHEJ = non-homologous end joining, ALS = amyotrophic lateral sclerosis). Values next to bars indicate the exact log10 Benjamini-Hochberg (B-H) adjusted p-values, with values below 10^-5^ rounded to 0. (B-F) Data reflects synthesis rates averaged from 3 biological replicates (see Fig. S3C for individual replicates).

This relationship was robust to alternative measures of AGA enrichment. A transcript-specific z-score representing the skew of arginine codon usage towards AGA compared to average genomic codon usage trends^2^ was also negatively correlated with the change in protein synthesis rate upon limitation for arginine, but not leucine (Fig. S3D). Binning genes by AGA content gave the same result; genes with high AGA frequency showed lower peptide levels upon arginine compared to leucine limitation, and genes with low AGA frequency had comparable synthesis rates levels following arginine or leucine limitation (Fig. 3D, S3E). These changes to synthesis rate affected steady state protein levels: higher AGA codon usage was associated with reduced protein abundance per mRNA upon limitation for arginine (Fig. S3F). Together, these data suggest that AGA codons are linked to lower protein synthesis rates and protein levels upon arginine limitation.

Upon leucine limitation in 293T cells, the reporter results suggest that UUA and UUG codons cause translation disruption in GCN2 KO cells (Fig. 1C,E). Consistent with this finding, we observed a stronger negative correlation between UUG usage and the change in protein synthesis rate upon leucine limitation in GCN2 KO cells compared to wild-type cells or to arginine limitation conditions (Fig. S3G). We did not observe a significant negative association between UUA usage and protein synthesis rate upon leucine limitation; translation disruption events may occur at clusters of UUA codons as in the reporter, but are not detectable across the entire transcriptome.

If protein synthesis rates are reduced directly by translation disruption upon amino acid limitation, this could result in depleted peptide representation at the C- versus the N-terminus (Fig. S3B). Indeed, the average fold change in peptide abundance decreased as a function of relative position along the length of a protein during arginine limitation (Fig. 3E); this positional bias was most evident following 24 hours of SILAC label incorporation and 5 total days of arginine limitation (Fig. S3H). Importantly, this effect was linked to AGA codon content: genes in the top quartile of AGA frequency exhibited pronounced depletion of C-terminal peptides during arginine limitation, whereas genes with zero or one AGA codon did not (Fig. 3F).

As expected, peptide positional effects were absent during leucine limitation in WT cells, consistent with the minimal translation disruption under these conditions (Fig. 3E). In contrast, GCN2 KO cells subjected to leucine limitation showed reduced abundance of C-terminal peptides, mirroring the reporter phenotype (Fig. 3E; Fig. 1C,E). Of note, extended leucine limitation led to reduced C-terminal peptides even in WT cells, suggesting that prolonged leucine limitation may eventually cause translation disruption on endogenous transcripts (Fig. S3H). Together, this analysis suggests that translation disruption can reduce endogenous protein production and increase the abundance of partially translated aberrant polypeptides in cells.

Finally, to identify endogenous transcripts most sensitive to translation disruption at AGA codons, we intersected genes with high AGA codon frequency and genes whose synthesis rates were reduced during arginine limitation in both WT and GCN2 KO cells. This yielded 118 genes enriched for functional categories including ribosomal proteins and mRNA processing factors, suggesting that translation disruption could have feedback effects on ribosome biogenesis and global mRNA maturation (Fig. 3G). Of note, given our measured proteome-wide correlations between AGA codon usage, protein synthesis rates, and C-terminal peptide depletion during arginine limitation, the effects of translation disruption likely extend to many genes.

### A genome-wide screen reveals the core regulatory network controlling translation disruption upon arginine limitation

To identify the mechanism and regulators of ribosome elongation disruption, we performed flow cytometry-based genome-wide CRISPRi/a screens in clonal K562 and 293T cells expressing Flag-YFP-4xAGA-DHFR and control Flag-mCherry-4xCGG-DHFR arginine codon reporters. Upon limitation for arginine, we sorted cells infected with a library of sgRNAs based on high or low YFP to mCherry “response” ratios and sequenced sgRNAs in each group. We then determined sgRNA enrichment in the high YFP versus low YFP response bins to calculate phenotype scores that specifically assessed translation disruption at AGA codons (Fig. 4A, Fig. S4A). Positive and negative phenotype scores indicate that the gene perturbation increased or decreased translation disruption or its product polypeptide levels, respectively. The hits (Fig. 4B-D) highlight a set of pathways that directly or indirectly regulate ribosome elongation at AGA codons during arginine limitation, converging on signaling responses that occur at stalled and/or collided ribosomes or in response to amino acid limitation including the mTORC1 and GCN2, ribotoxic stress response (RSR), and ribosome splitting quality control trigger (RQC-T) and quality control (RQC) pathways (Fig. 4E).

**Figure 4:**
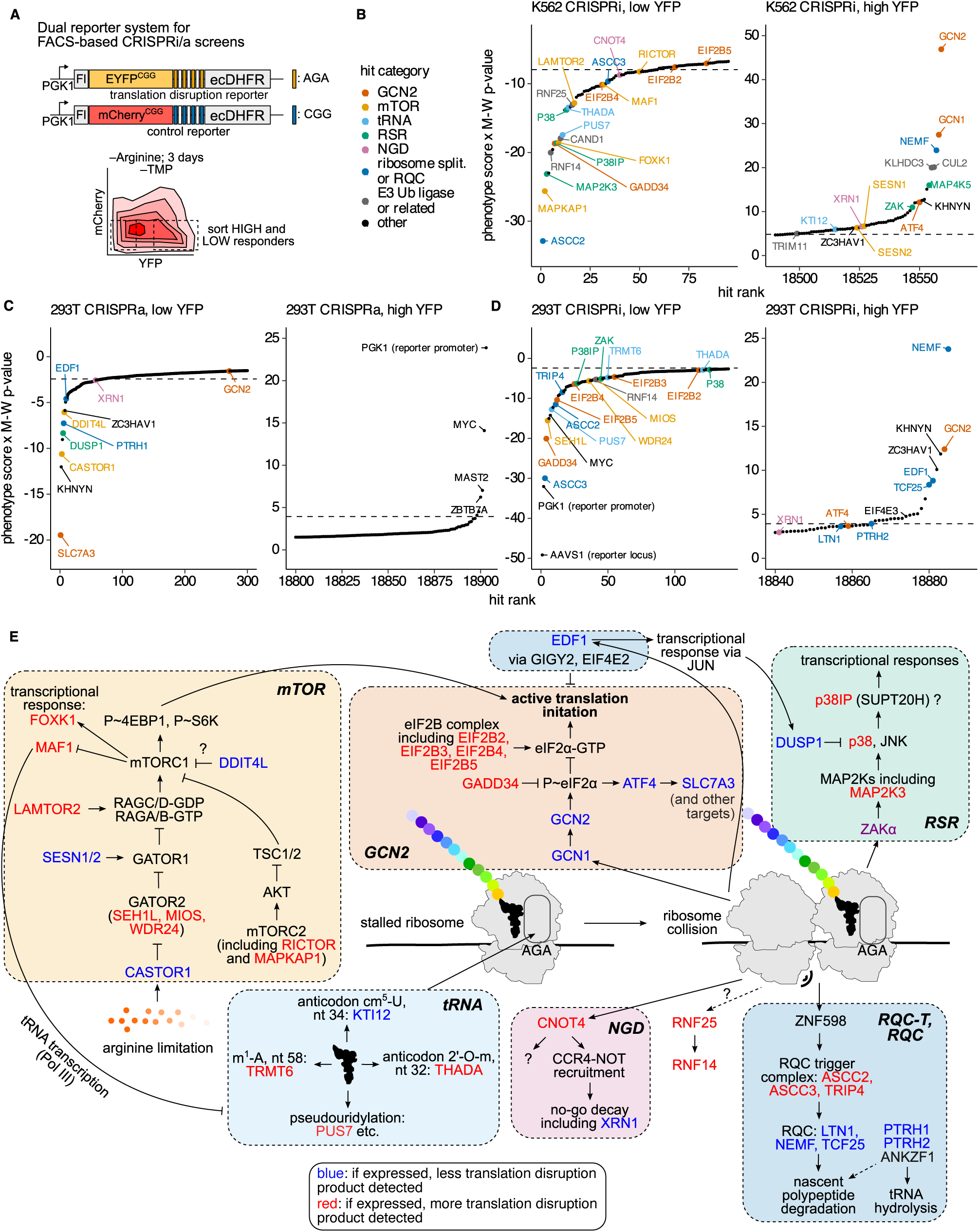
A genome-wide screen reveals the core regulatory network controlling translation disruption upon arginine limitation. (A) A dual reporter system with a Flag-mCherry^CGG^-4xCGG-DHFR^CGG^ (internal control, no translation disruption) and a Flag-YFP^CGG^-4xAGA-DHFR^CGG^ was integrated into the AAVS1 locus in cells expressing CRISPRi or CRISPRa machinery (see Methods) to query the effect of a genome-wide library of CRISPRi/a sgRNAs on specific translation disruption signal. Flow cytometry was used to sort high and low responding populations. (B-D) Total score (phenotype score * Mann-Whitney (M-W) p-value, see Methods) versus hit rank for highest (high YFP) and lowest (low YFP) scoring genes in K562 CRISPRi (B), 293T CRISPRa (C), and 293T CRISPRi (D) screens. Hits are labeled and colored by associated function. Dashed line marks score cut-off where FDR = 0.25. (E) Diagram connecting major screen hit pathways. Hits are colored by direction of phenotype; blue gene labels reduce translation disruption product accumulation, red gene labels increase translation disruption product accumulation.

Several RNA modifying enzymes were also hits in the screen. These included two genes that encode factors that modify the tRNA anticodon loop: the methyltransferase THADA, which modifies position 32 with 2’-O-ribose, whose expression stimulated translation disruption (Fig. 4B,D, Fig. S4B,C); and KTI12, a component of the Elongator complex that modifies Arg-tRNA^UCU^ with mcm^5^ at U34 to increase AGA decoding efficiency^66–69^, whose expression reduced translation disruption as expected (Fig. 4B). The tRNA m^1^A methyltransferase TRMT6 and the RNA pseudouridylase PUS7 also promoted translation disruption (Fig. 4B,D). While only KTI12 has a previously demonstrated link to Arg-tRNA^UCU^, these results suggest that a range of tRNA and mRNA modifications can influence ribosome elongation during arginine limitation.

### Stress response signaling pathways regulate arginine-limitation induced translation disruption

The mTORC1 and GCN2 pathways, whose responses to amino acid limitation repress translation initiation, are known regulators of ribosome stalling^2,42,70,71^. Several hits confirmed this and specified the nodes of these pathways that regulate translation disruption. For example, the integrated stress response kinase GCN2 and its activator GCN1, which respond to arginine limitation through eIF2α phosphorylation and induction of the transcription factor ATF4, all reduced translation disruption (Fig. 4B,D; Fig. S4B,C). Conversely, the phosphatase GADD34, which attenuates eIF2α phosphorylation, and the eIF2B complex, which activates global protein synthesis via eIF2α, promoted translation disruption (Fig. 4B,D; Fig. S4D). mTORC1 kinase activity, which is partially inhibited upon arginine limitation^2,72^, had the opposite effect as expected. mTORC1 activators including the GATOR2 complex, LAMTOR2, and components of mTORC2 stimulated translation disruption (Fig. 4B,D, Fig. S4B,C), whereas the leucine and arginine sensors/mTORC1 inhibitors SESN1/2^73–77^ and CASTOR1^78,79^ protected against translation disruption (Fig. 4B,C). We confirmed that the mTOR inhibitor Torin1 also reduced translation disruption (Fig. S4E,F). Regulation of translation disruption by the GCN2 and mTORC1 responses to amino acid limitation can likely be attributed to reduced translation initiation, as higher initiation rates would both consume limited arginine and promote ribosome collisions, which directly trigger translation disruption through the RQC pathway^20^ and frameshifting^80^. In support of this possibility, EDF1, a ribosome collision sensor that reduces translation initiation *in cis*^81,82^, lessened translation disruption (Fig. 4C,D).

Ribosome collisions also activate the ribotoxic stress response (RSR) signaling cascade involving MAP3K ZAKα, MAP2Ks, p38 and JNK^42^. p38 and MAP2K3 increased translation disruption (Fig. 4B,D; Fig. S4B,C), and the p38 phosphatase DUSP1^83–85^ reduced translation disruption (Fig. 4C). ZAKα had divergent effects on translation disruption in the two cell models screened (Fig. 4B,D), perhaps due to its proposed position upstream of both the GCN2 and p38 responses^42^, which had opposite effects on translation disruption (Fig. 4E).

To confirm that arginine limitation induces the RSR, we measured levels of phosphorylated p38 and found that they increased over time, especially in GCN2 KO cells, though the response was weaker than that induced by exposure to an elongation inhibitor or UV irradiation (Fig. S4B,G,H), controls known to induce ribosome collisions^42^. We did not detect a JNK phosphorylation response to arginine limitation (Fig. S4G). p38 phosphorylation was higher during limitation for arginine than leucine (Fig. S4G), consistent with more translation disruption following arginine limitation (Fig. 1B,C,E). We generated ZAKα KO and ZAKα/GCN2 double KO 293T cells (Fig. S4I) to verify that p38 phosphorylation was ZAKα-dependent (Fig. S4J-K). MAP2K3 knockdown also prevented p38 phosphorylation during arginine limitation in K562 cells, suggesting that it represents the intermediate kinase between ZAKα and p38 in these cells (Fig. S4B).

We used a p38 kinase inhibitor to confirm that loss of p38 activity reduces translation disruption upon arginine limitation (Fig. S4L,M). The inhibitor increased p38 phosphorylation during arginine limitation (Fig. S4H), suggesting negative feedback downstream of the p38 response. p38 inhibition also reduced reporter mRNA accumulation (Fig. S4N), suggesting the p38 pathway regulates reporter transcription upon arginine limitation, as previously proposed for virally-integrated transgenes^58,86^ (see Fig. S1U,Z). However, if reporter mRNA accumulation is an *in trans* consequence of global ribosome collision signaling^58^, any perturbation that suppresses ribosome collisions should indirectly reduce reporter mRNA accumulation. In line with this possibility, we confirmed that knockdown of LAMTOR2, which reduces translation disruption by dampening mTORC1 signaling, similarly prevented reporter mRNA accumulation during arginine limitation (Fig. S4O). These data are consistent with a model in which the RSR through p38 stimulates either translation disruption or the downstream reactivation of viral expression elements in the genome.

### The ribosome splitting and quality control pathways resolve stalled ribosomes during arginine limitation

The ribosome quality control (RQC) trigger (RQC-T) and RQC pathways respectively execute ribosome splitting and nascent polypeptide degradation downstream of collisions caused by stalling on damaged or defective mRNAs^22,23,87^. Multiple screen hits suggested that RQC-T and RQC also act on intact mRNAs with stalled, collided ribosomes during arginine limitation. First, loss of any component of the RQC-T complex (ASCC2/ASCC3/TRIP4) reduced arginine-limited translation disruption (Fig. 4B,D; Fig. 5A; Fig. S5A). Knockdown of LTN1, NEMF, and TCF25, the RQC proteins that target nascent polypeptides on split ribosomes for degradation^45,88–93^, and the upstream peptidyl-tRNA hydrolases PTRH1/2^93^, increased translation disruption product levels (Fig. 4B-D; Fig. 5B,C), confirming that the aberrant polypeptide products are targeted by RQC.

**Figure 5:**
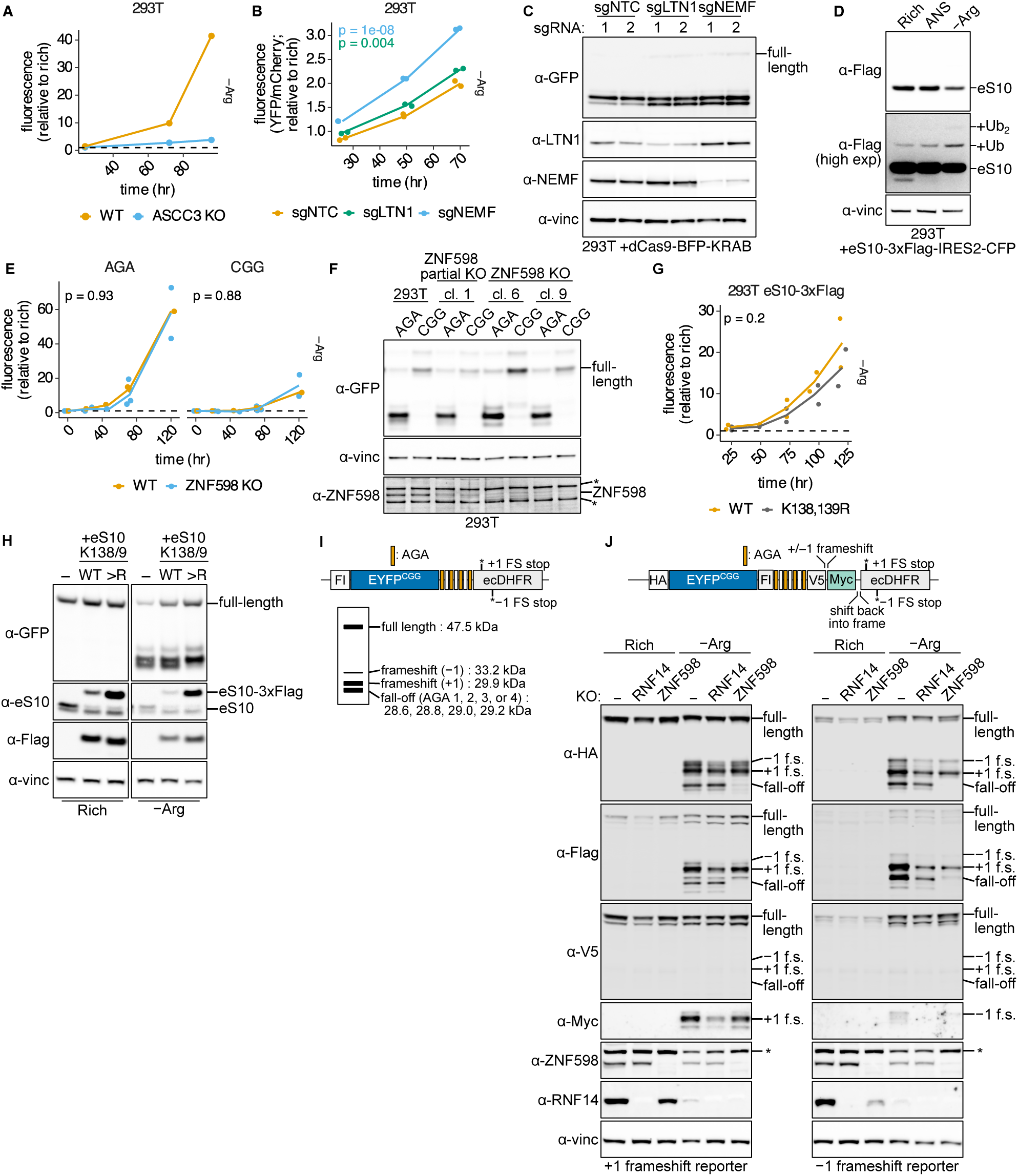
The ribosome splitting and quality control pathways resolve stalled ribosomes during arginine limitation. (A) Flow cytometry to assess translation disruption product accumulation with or without ASCC3 KO upon limitation for arginine in 293T cells expressing the Flag-YFP^CGG^-4xAGA-DHFR^CGG^ reporter. (B) Flow cytometry to assess dual color translation disruption reporter signal accumulation upon limitation for arginine (Flag-YFP^CGG^-4xAGA-DHFR^CGG^ over control Flag-mCherry^CGG^-4xCGG-DHFR^CGG^) with or without LTN1 or NEMF knockdown using CRISPRi in 293T cells. (C) Western blot to assess translation disruption upon LTN1 or NEMF knockdown with 2 sgRNAs using CRISPRi and arginine limitation for 3 days in 293T cells expressing the dual color translation disruption reporter integrated at the AASV1 locus. (D) Western blot to assess eS10-3xFlag ubiquitination upon treatment with 0.1 μg/mL anisomycin (ANS) for 15 minutes or arginine limitation for 3 days in 293T cells expressing eS10-3xFlag-IRES2-CFP. +Ub or +Ub_2_ labels mark mono- and di-ubiquitination. (E,F) Flow cytometry (E) or western blot (F) to assess translation disruption product accumulation with or without ZNF598 KO across 3 clones (“cl.”) upon limitation for arginine over time (E) or for 5 days (F) in 293T cells expressing the Flag-YFP^CGG^-4xAGA-DHFR^CGG^ (“AGA”) or Flag-YFP^CGG^-4xCGG-DHFR^CGG^ (“CGG”) reporters as indicated. (G,H) Flow cytometry (G) or western blot (H) to assess translation disruption product accumulation in 293T cells expressing WT or K138,139R (“>R”) mutant eS10-3xFlag-IRES2-CFP upon arginine limitation over time (G) or for 5 days (H) in 293T cells expressing the HA-YFP^CGG^-4xAGA-DHFR^CGG^ reporter. (I) Flag-YFP^CGG^-4xAGA-DHFR^CGG^ reporter and example western blot schematic identifying major translation disruption product bands. (J) Western blot to assess translation disruption including fall-off and +/−1 frameshifting (f.s.) in wild-type, RNF14 KO, or ZNF598 KO 293T cells expressing the illustrated multi-tag frameshifting reporters upon limitation for arginine and treatment with 10 μM TMP for 3 days. (B,E,G) ANOVA used to determine significance with pairwise differences assessed using estimated marginal means with a Tukey correction for multiple testing where applicable; p-values are shown on the plot. In B, pairwise p-values represent sgNTC vs sgGuide comparison and are colored according to guide. (A,E,G) Dashed lines at y = 1. (C,F,H,J) “*” marks a non-specific band on ZNF598 blot. Full-length reporter product is indicated (α-GFP antibody used to detect YFP, α-vinc = α-vinculin).

Collided ribosomes can trigger substrate mRNA decay via the no-go decay (NGD) pathway^94,95^ and the NGD exonuclease XRN1 and E3 Ubiquitin ligase CNOT4^96–98^ scored as weak hits in the screens (Fig. 4B-D). However, targeted knockdown of these genes did not affect reporter mRNA level or overall translation disruption product accumulation (Fig. S5B,C). Knockdown of N4BP2, whose yeast homolog Cue2 catalyzes endonucleolytic cleavage between collided ribosomes in the absence of ribosome splitting^99^, also had little effect (Fig. S5B,C). Thus, mRNA decay is not strongly coupled to translation disruption during arginine limitation.

Unexpectedly, the E3 ubiquitin ligase ZNF598 that senses ribosome collisions on damaged or defective mRNAs and ubiquitinates eS10/uS10 (RPS10/20) to trigger ribosome splitting^17–19,100^ was not a hit in the screens (Fig. 4B-D). Since we detected an increase in ubiquitinated eS10 upon arginine limitation (Fig. 5D), we both generated ZNF598 knockout cells (Fig. S5D) and ectopically expressed the eS10 K138,139R ubiquitin site mutant, which largely replaces endogenous eS10 to block eS10 ubiquitination^17–19^, to assess their role in translation disruption. ZNF598 KO and loss of eS10 ubiquitination minimally affected total levels of translation disruption upon arginine limitation (Fig. 5E-H). eS10 levels were reduced by arginine limitation and the K138,139R mutant increased eS10 levels (Fig. 5H), consistent with its ubiquitination leading to 40S degradation^101–107^. However, loss of ZNF598 or eS10 ubiquitination clearly altered the mode of translation disruption, as assessed by aberrant polypeptide product mobility. Accumulation of a shorter polypeptide depended on eS10 ubiquitination (Fig. 5F,H), whereas loss of RQC factors LTN1/NEMF stabilized it (Fig. 5C). Thus, this shorter translation disruption product represents the ribosome “fall-off” product caused by ribosome splitting at the AGA codon linker (Fig. 5I). The prevalent longer translation disruption product independent of RQC activity likely represents the result of +1 frameshifting at AGA codons, which would lead to premature stop codon recognition after 12 amino acids (Fig. 5I). Similarly, a slightly longer product corresponds to a −1 frameshifting event (Fig. 5I). We confirmed that AGA codons lead to +1, and less so, to −1 frameshifting products upon arginine limitation using a modified version of the translation disruption reporter with a +1 or −1 Myc tag (Fig. 5J). This Myc tag reporter construct, which increased resolution of the fall-off and frameshifting products, confirmed a requirement of ZNF598 for the shorter polypeptide product of ribosome fall-off directly at AGA codons (Fig. 5J). These data suggest that ZNF598 controls the mode, but not overall amount, of ribosome elongation failure during arginine limitation, and that the RQC-T and RQC pathways are key downstream steps in ribosome fall-off.

### The E3 ubiquitin ligase RNF14 promotes translation disruption at arginine-limited ribosomes

The E3 ligase RNF14 and its upstream activating E3 ligase RNF25 are implicated in degradation of ribosomal A-site obstructions^36,37^ and drug-trapped eEF1A^34^ and eRF1^35^ translation factors. Knockdown of these ubiquitin ligases reduced translation disruption in the screen (Fig. 4B,D). We confirmed RNF14 knockout reduced the accumulation of both frameshifting and fall-off translation disruption products upon arginine limitation (Fig. 5J, Fig. 6A,B). Importantly, neither eEF1A nor eRF1 were degraded during arginine limitation (Fig. S6A,B), suggesting a novel role for RNF14 in the absence of ribosomal A-site obstructions.

**Figure 6:**
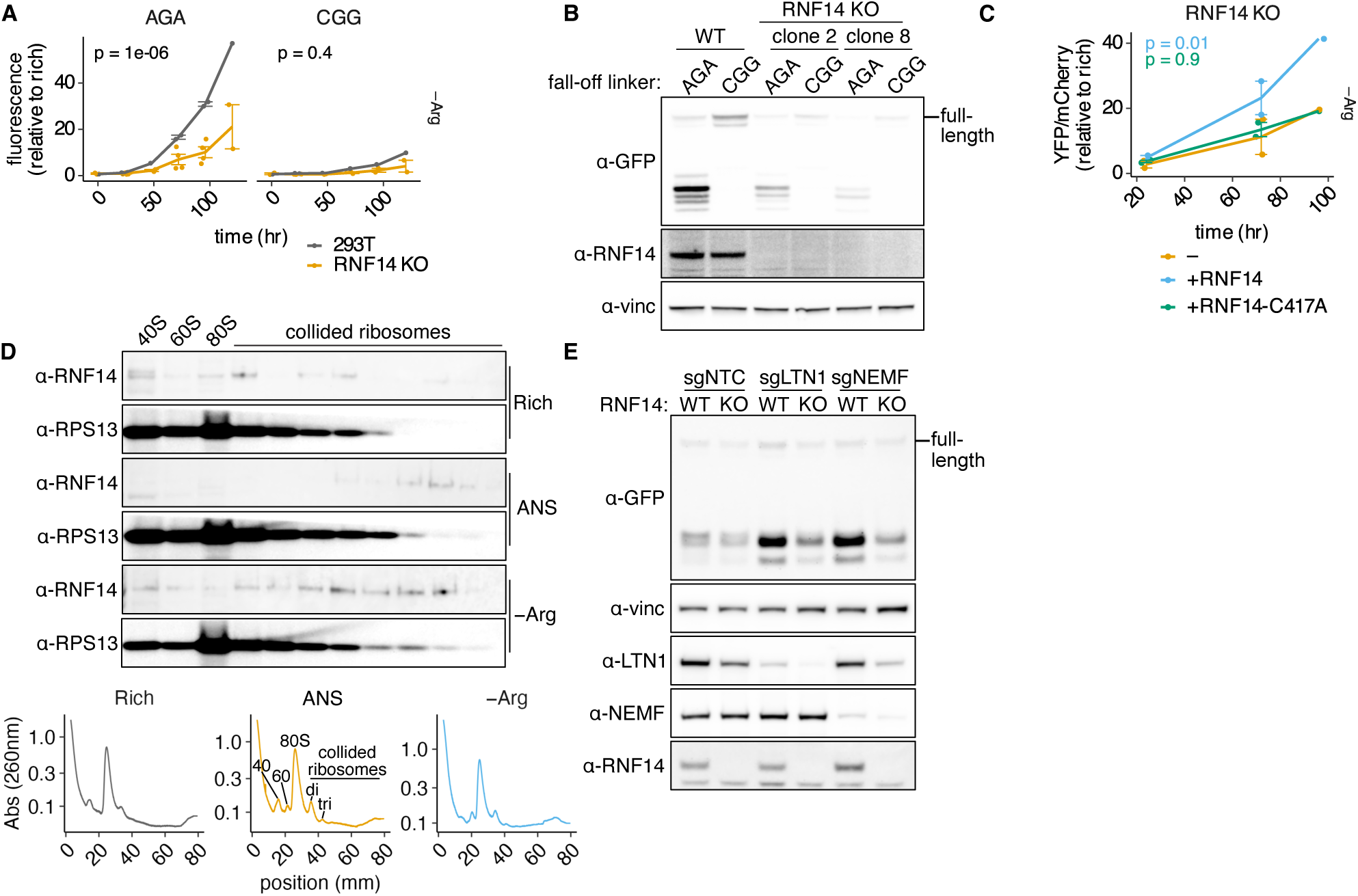
The E3 ubiquitin ligase RNF14 promotes translation disruption at arginine-limited ribosomes. (A-B) Flow cytometry (A) or western blot (B) to assess translation disruption product accumulation with or without RNF14 KO across 2 replicate clones in 293T cells expressing the Flag-YFP^CGG^-4xAGA-DHFR^CGG^ (“AGA”) or Flag-YFP^CGG^-4xCGG-DHFR^CGG^ (“CGG”) reporters over time upon arginine limitation (A) or after 5 days of arginine limitation (B). (C) Flow cytometry to assess translation disruption product accumulation in RNF14 KO 293T cells expressing nothing, wild-type, or C417A mutant V5-tagged RNF14 and the Flag-YFP^CGG^-4xAGA-DHFR^CGG^ reporter upon arginine limitation. (D) Western blot on sucrose density gradient fractions from MNase-treated lysate to determine RNF14 association with collided polyribosomes and ribosome subunits in rich conditions, upon arginine limitation for 5 days, or after 15 minutes of treatment with 0.1 μg/mL anisomycin (ANS). 260 nm absorbance profiles collected during fractionation are shown below the blots; peaks are labeled with associated ribosome fraction. (E) Western blot to assess translation disruption product accumulation after 2 days of arginine limitation in wild-type or RNF14 KO 293T cells with or without knockdown using a non-targeting control (NTC), or guide RNAs targeting LTN1 or NEMF as indicated. (A,C) Error bars represent the average of 2-4 biological replicates; ANOVA used to determine significance with pairwise differences assessed using estimated marginal means with a Tukey correction for multiple testing where applicable; p-values are shown on the plot. P-values represent comparison to wild-type controls and in C are colored according to RNF14 construct. (B, E) Full-length reporter product is indicated (α-GFP antibody used to detect YFP, α-vinc = α-vinculin)

While ectopic RNF14 expression increased translation disruption in RNF14 KO cells, a catalytic site mutant (C417A) of RNF14 did not, confirming that ubiquitin ligase activity is required to stimulate translation disruption (Fig. 6C). RNF14 levels fell progressively during arginine limitation, likely reflecting auto-degradation due to sustained activation^34,108^ (Fig. S6B). Confirming its localization to stalled ribosomes, RNF14 was associated with nuclease-resistant collided ribosomes after sucrose density gradient fractionation, though was also present in these fractions in nutrient rich conditions (Fig. 6D). The collision sensor GCN1 was previously found to activate RNF14^34–37;^ however, we found in our screen that GCN1 expression prevented translation disruption (Fig. 4B), likely due to its dominant role as a mediator of GCN2 activation. We checked whether RNF14 might degrade RQC factors, as several were previously reported to be RNF14 targets^37^. Upon arginine limitation, RNF14 loss decreased LTN1 levels and did not affect NEMF levels, inconsistent with both a role in degrading RQC factors and the direction of its effect on translation disruption product accumulation (Fig. S6B).

We next explored the consequences of RNF14 loss for translation more generally. When we stabilized the translation disruption reporter degron to assess full-length reporter production, RNF14 loss did not increase read-through past the translation disruption site as ZNF598 loss does for other RQC reporters^19,100^ (Fig. S6C). RNF14 loss also did not measurably increase ribosome collision signaling through p38 phosphorylation or alter global translation rate during growth in nutrient rich conditions or arginine limitation (Fig. S6C,D).

To test the hypothesis that RNF14 acts upstream of translation disruption during arginine limitation, we performed a pathway epistasis analysis. While LTN1 or NEMF loss increased stable translation disruption product levels in WT cells as expected, their loss had markedly less impact on translation disruption product levels in RNF14 KO cells (Fig. 6E, Fig. S6E), suggesting that nascent polypeptide quality control pathway flux is reduced in the absence of RNF14 activity. As RNF14 loss also reduces the frameshifted polypeptide translation disruption product (Fig 5J), it likely acts upstream of RQC initiation. Interestingly, RNF14 loss did not strongly affect translation disruption reporter activity at affected threonine or leucine codons, highlighting a potentially unique mechanistic link between arginine limitation and RNF14 activity (Fig. S6F).

## DISCUSSION

Ribosome stalling due to charged tRNA limitation has been linked to diverse normal and disease physiology including cancer^5,9,27,50^, neurodegeneration^8^, and biofilm formation^7^ and can cause premature translation disruption. We find translation disruption varies widely based on the specific amino acid limitation conditions and across cell lines. Depletion of several amino acids, including lysine, phenylalanine, histidine, and methionine, caused substantial charged tRNA loss but little to no translation disruption reporter signal (Fig. S2A). In these cases, ribosome stalling may be resolved through mechanisms such amino acid misincorporation. We note that although we attempted to identify robustly translated synonymous codons for encoding YFP, ruling out false negatives caused by disruption of protein synthesis earlier in YFP remains challenging, particularly for amino acids with few synonymous codons or cognate tRNAs. Nevertheless, arginine limitation appears to be a uniquely consistent strong trigger for translation disruption. The kinetics of translation disruption hours after arginine limitation are somewhat surprising as elongation inhibitors can cause collisions within minutes^42^. The data support a model wherein charged Arg-tRNA^UCU^ levels gradually decline, leading to a gradual onset of ribosome stalling, collisions and translation disruption that impacts the proteome.

Synonymous codon-specific translation disruption can be explained by differences in isoacceptor tRNA charging, raising the question of why isoacceptors charged by the same synthetase respond differently to amino acid limitation. AGA is the most abundant arginine codon and Arg-tRNA^UCU^ is the rarest tRNA, creating a supply-demand imbalance that has been proposed to explain a similar phenomenon in *E. coli*^43,44^. Other models have been proposed, including differential tRNA competition for ribosome recruitment^109^ and tRNA levels as a predictor of ribosome frameshifting^4^. In mouse models loss and gain of an Arg-tRNA^UCU^ tRNA gene have been linked to neurodegeneration^8^ and cancer progression^110^, respectively, suggesting that a critical set point has evolved to make translation at this codon responsive to metabolic state, though whether and how this affects cell function remains unclear.

The mechanisms of stalled ribosome quality control (RQC) have been well-studied in the context of mRNA defects such as rare codons or premature polyA tracts, and less so at ribosomes stalled due to amino acid limitation on intact mRNAs. Our findings add to the list of endogenous targets for the RQC pathway. The ribosome splitting complex RQC-T is a strong regulator of ribosome fall-off during arginine limitation, as is downstream peptide targeting by the RQC pathway. The peptide targeting E3 ligase LTN1 was a weaker hit in the screen, consistent with previous reports that it does not efficiently ubiquitinate the nascent GFP polypeptide due its secondary structure^88^ and that CAT-tailing by NEMF can act as a ubiquitin-independent route for nascent peptide targeting^111^. The RQC collision sensor ZNF598 was not a regulator of overall translation disruption during arginine limitation, instead shifting the mode of elongating ribosome failure from RQC-mediated ribosome fall-off to +1 frameshifting, consistent with previous reports that ZNF598 loss increases frameshifting^112^. This contrasts with observations that ZNF598 loss promotes read-through on polyA tracts^19,100^, and we speculate that read-through is ultimately limited by charged tRNA concentrations during amino acid limitation. Further, loss of the downstream RQC-T complex reduced overall translation disruption, suggesting that ZNF598-mediated ribosome ubiquitination may reduce the probability of frameshifting.

We did not find evidence for a strong link between ribosomes stalled due to arginine limitation and mRNA decay pathways. The non-stop decay (NSD)-initiating factor Pelota, and its downstream partners HBS1L, ABCE1, which release ribosomes with empty A-sites stalled at 3’ transcript ends^95^ to trigger mRNA decay^113^, were not hits in our screens; this is likely explained by the reported steric incompatibility of A-site mRNA with Pelota binding^114^ and its consequent lack of activity on mRNAs with more than 9 nucleotides following the stall site^115^.

RNF14 is an E3 ubiquitin ligase with a proposed role in stalled ribosome resolution, clearing obstructions in the ribosome A-site caused by drugs that trap translation factors in the ribosome or following the induction of RNA-protein crosslinks^34–37^. We identify an additional role for RNF14 affecting tRNA-limited ribosomes where the A-site is unobstructed. RNF14 has been reported to be activated by the collision sensor GCN1^34–37^, and our results place it upstream of both RQC-mediated ribosome fall-off and ribosome frameshifting. Although we did not detect a difference in ribosome collision signaling or global translation rates upon RNF14 loss, an attractive model is that RNF14 affects ribosome collision frequency, which stimulates both fall-off and frameshifting. Future work to determine the targets of RNF14 and their mechanistic activity at the stalled ribosome will reveal its role in translation disruption during amino acid limitation.

## MATERIALS AND METHODS

### Cell culture

Cells were grown at 37°C and 5% CO_2_ in humidified incubators. Cell line identity and quality was confirmed in early passage stocks used as “parents” for all cell line generation through verification by ATCC and mycoplasma testing using the MycoAlert mycoplasma detection kit (Lonza LT07-318). HEK293T (CRL-3216), HCT 116 (CCL-247), HeLa (CCL-2), PANC-1 (CRL-1469), MiaPaCa (CRL-1420), MDA-MB-231 (HTB- 26), and MS-1579 cells (primary mouse breast cancer cells^52^) were cultured in DMEM without pyruvate +10% heat inactivated FBS, and K562 (CCL-243) cells were cultured in RPMI +10% FBS. For CRISPRi/a screens only, cells were culture with penicillin and streptomycin at final concentrations of 100 U/mL and 100 μg/mL, respectively (Corning 30–002–CI).

For amino acid limitation, cells were plated in 6 well or 6 cm plates in standard culturing medium the day before amino acid withdrawal such that they would be 50-80% confluent the next day (or at the time of harvest, for Rich medium controls). They were washed once in excess PBS after aspiration, and medium was replaced with DMEM lacking the amino acid of interest +10% heat inactivated dialyzed FBS (see Preparation of amino acid dropout medium). The maximum recommended volume of medium per well or plate was used to minimize depletion of other nutrients across extended amino acid limitation experiments. Cell density impacted the severity of amino acid limitation, with confluency protective and sparse plating more sensitive, and thus translation disruption reporter responses were performed on a defined number of cells to ensure reproducible responses; for adherent cell lines, 500,000 cells were plated in a 6 well plate well or equivalent cell number to surface area ratio in larger plates. For suspension cell experiments, cell density was initially 0.2 million/mL.

### Preparation of amino acid dropout medium

DMEM lacking every amino acid was prepared from a low-glucose powdered DMEM base lacking amino acids and pyruvate (US Bio 9800-13). 8.32 g DMEM base was combined with 3.5 g glucose and 3.7 g sodium bicarbonate per liter. Custom amino acid mixes containing groups of DMEM amino acids were combined by weighing out enough of each amino acid for 20 L and combining powders to homogeneity using a coffee grinder (Hamilton Beach), stored in 50 mL Falcon tubes at −20°C, and supplemented to the appropriate weight per liter. Homemade amino acid limitation medium was filtered using a 0.45 μM vacuum flask with 10% dialyzed FBS. Some lots of dialyzed FBS displayed protein aggregation after heat inactivation and were spun at 5000 RPM in a tabletop Beckman Coulter centrifuge for 10 minutes to pellet protein before filtration.

### Cloning reporter variant library

Translation disruption reporters were cloned into the lentiviral donor vector pLJM1 containing an N-terminal Flag tag and C terminal ecDHFR degron tag from ref ^2^ (adapted from ref ^116^). For each amino acid: gene blocks containing YFP codon variants were ordered from IDT. SalI and EcoRV digestion allowed Gibson assembly cloning of codon variants generated pLJM1-Flag-YFP^Codon^ ^Variant^-DHFR (Figure 1A, step 1). YFP synthesis rate was compared after addition of 10 μM TMP to determine the optimally translated codon for YFP (“safe”). Then, to create translation disruption reporter library, SalI and EcoRI digestion allowed Gibson assembly of YFP^safe^ and DHFR amplified using primer overhangs to add the 4x codon variant linkers (Figure 1A, step 2).

### Lentivirus preparation and

#### sgRNA vectors

HEK293T lenti-X cells (Takara) were transfected with a lentiviral donor plasmid, psPAX2 (Addgene #12260) and pCMV-VSV-G (Addgene #8454) in a 2:2:1 molar ratio using Lipofectamine 3000 (Invitrogen) according to the manufacturer’s instructions. The culture medium was collected at 24–30 h and 48–54 h post-transfection, pooled, centrifuged at 1,000 x *g* for 10 min, and filtered through a 0.45 μm low-protein-binding membrane. Lentivirus-containing supernatants were used immediately or stored at −80 °C. All lentiviral transductions of target cell lines were performed in the presence of 8 mg/mL polybrene (Sigma).

### Translation disruption reporters/cDNA overexpression

HEK293T lenti-X cells (Takara) were transfected with a lentiviral donor plasmid, pMDLg/RRE (Addgene #12251), pMD2.G (Addgene #12259), and pRSV-Rev (Addgene #12253) in a 4:2:2:1 molar ratio using Lipofectamine 3000 (Invitrogen) according to the manufacturer’s instructions. The culture medium was collected at 24–30 h and 48–72 h post-transfection, pooled, and filtered through a 0.45 μm low-protein-binding membrane. Lentivirus-containing supernatants were used immediately or stored at −80 °C. Transduction was performed by infecting ∼200,000 cells with 500 μL viral supernatant medium, changing the medium after 12-24 hours, and performing selection with puromycin after 48 hours (2 μg/ml for HEK293T and K562 cells, 1 μg/ml for all other cell lines).

### Cell line generation

#### Generation of CRISPRi/a parental cell lines expressing translation disruption reporters

To generate a K562 cell line stably expressing dCas9-KRAB and the translation disruption reporters, K562 parental CRISPRi monoclonal cells from ref ^117^ were transfected with reporter plasmids mCherry^CGG^-4xCGG-ecDHFR^CGG^ and YFP^CGG^-4xAGG-DHFR^CGG^, and pX330-AAVS1 (1:1:1 w/w/w) using Lipofectamine 3000 (Invitrogen) according to the manufacturer’s instructions. Starting at 5 days post-transfection, cells were selected using blasticidin at 10 mg/mL for 10 days. Cells were cultured for 2 days in RPMI +10 μM TMP and single cells expressing both mCherry and YFP were selected by fluorescence-activated cell sorting (FACS) (BD FACS Aria II). Monoclonal cultures were expanded and the best performing clone was selected by analyzing the reporter response in −Arg medium (see Preparation of amino acid dropout medium).

To generate a HEK293T cell line stably expressing dCas9-KRAB and dual color translation disruption reporters, HEK293T +Flag-YFP^CGG^-4xAGA-DHFR^CGG^ +Flag-mCherry^CGG^-4xCGG-ecDHFR^CGG^ clone 9 cells were infected with lentiviral particles produced using vector pMH0001 (Addgene #85969). A polyclonal population of cells expressing dCas9-KRAB was generated by FACS (BD FACS Aria II) by gating on the top half of BFP-positive cells. In a final step, a pure population of cells expressing both translation disruption reporters were selected by FACS gated on mCherry- and YFP-positive cells after growth for 2 days in 10 μM TMP.

To generate a HEK293T cell line stably expressing the CRISPRa machinery and translation disruption reporters, HEK293T +Flag-YFP^CGG^-4xAGA-DHFR^CGG^ +Flag-mCherry^CGG^-4xCGG-ecDHFR^CGG^ clone 9 cells were infected with lentiviral particles produced using vector HRdSV40-dCas9-10xGCN4_v4-P2A-BFP (Addgene #60903). BFP^+^ cells were selected by FACS and then infected with lentiviral particles produced using vector pHRdSV40-scFv-GCN4-sfGFP-VP64-GB1-NLS (Addgene #60904). BFP and GFP double positive single cells were selected by FACS into single wells of a 96-well plate. Monoclonal cultures were expanded and the best performing clone was selected by analyzing the reporter response in −Arg medium, and by testing the performance of the CRISPRa system using 3 test sgRNAs and RT-qPCR. In a final step, cells expressing both translation disruption reporters were selected by FACS gated on mCherry- and YFP-positive cells after culture for 2 days in 10 μM TMP.

#### Generation of knockout cell lines

Knockout cell lines were prepared as described in ref ^2^ with the following modifications. Two sgRNAs were designed in introns flanking the first coding exon that was not a multiple of 3 in length and cloned into the targeting plasmids pU6-(BbsI)_CBh-Cas9-T2A-BFP (#64323) and pSpCas9(BB)-2A-GFP (Addgene #48138). Knockout clones were verified by genomic DNA PCR using cell pellets lysed in QuickExtract buffer (Lucigen) according to the manufacturer’s instructions, using primers flanking the sgRNA binding sites; and/or by western blotting.

### Generation of knockdown and overexpression cell lines

CRISPRi/a cell lines targeting individual genes were prepared in the CRISPRi/a parental cell lines expressing translation disruption reporters, and changes in expression level were quantified, as previously reported ref ^117^.

### Analytical flow cytometry

For analytical flow cytometry, cells were harvested form a 6 well plate well by PBS wash and trypsinization, pelleted at 200 x *g* for 3 minutes, resuspended in PBS, pelleted at 200 x *g* for 3 minutes, resuspended in 250 μL PBS, and passed through a mesh-top flow cytometry tube. After adjusting laser voltages based on unstarved controls and gating to isolate single cells, 200,000 events were collected. Raw data was analyzed in R using the flowCore package to read .fcs files, gated, and the mean of 200,000 events was calculated. Reporter fluorescence in amino acid limited conditions was normalized to fluorescence in nutrient rich conditions to correct for any effects of codon usage on baseline expression levels.

To evaluate the effect of different genetic perturbations on reporter signal quantified by flow cytometry, we used estimated marginal means from a fitted ANOVA model and performed pairwise comparisons using the emmeans() package in R. P-values were calculated with adjustment for multiple comparisons using Tukey’s honestly significant difference (HSD) method.

### RTqPCR

Plates or pellets of cells were washed in PBS and flash frozen in liquid nitrogen. Monolayers or pellets were lysed in RNA lysis buffer (Zymo) and total RNA purified using the Zymo RNA miniprep kit including the DNaseI digestion step to remove genomic DNA. 500 ng RNA was reversed transcribed to cDNA using SuperScript IV (NEB) with an oligo-dT25 primer. cDNA was diluted 1:5 in water and qPCR performed using the PowerUp SYBR Green Master Mix (Applied Biosystems) according to manufacturer’s instructions for reaction components, cycle temperatures and times. Samples were run on a Roche Lightcycler 480 II.

### Western blotting

Plates or pellets of cells (∼250,000-2 million cells) were washed in PBS and flash frozen in liquid nitrogen. Monolayers were scraped into, or pellets were lysed in, 1X RIPA buffer on ice (60 μL per 6 well plate well or equivalent volume per surface area ratio in larger plates). Lysate was clarified at 21,000 x *g* at 4°C for 10 minutes. Absolute protein concentration was measured using a BCA assay (Pierce) and albumin standard curve. 5-25 μg of protein in 1x LDS sample buffer (Novex) with 50 mM DTT was loaded onto 4-12% Bolt Bis-Tris precast gels (Novex) and run at 125V for 90 minutes. Gels were rinsed in 1X Tris-Glycine transfer buffer with 20% methanol and semi-dry transfer was performed to 0.45 μm nitrocellulose at 18V for 60 minutes (Bio-Rad Trans-Blot SD). Membranes were blocked with 5% ultra pure BSA (CST) or nonfat milk. Primary antibody incubation was performed overnight at 4°C followed by 3x 5 minute 1x TBST washes, 1-5 hour secondary goat anti-mouse or anti-rabbit IgG at 1:5000 (CST), 3x 5 minute 1x TBST washes, and then developed using Western Lighting ECL reagent (Pierce) and imaged using the auto-expose setting on an ImageQuant LAS-4000 cooled camera system (GE). Membranes were stripped with Restore Western blot stripping buffer (Thermo Fisher) for 15 minutes followed by 1-5 hours shaking at 37°C with 30% H_2_O_2_ before re-blocking.

### Puromycin pulse to measure protein synthesis rate

1 million HEK293T cells were plated in a 6 cm plate. The next day, cells were washed and treated with arginine limitation medium. After 3 days of arginine limitation or 30 minutes of treatment with 10 μg/mL cycloheximide as a control, puromycin was spiked into the culture to 10 μg/mL. Before spike in, or after 30, 60, 120, or 240 seconds, cells were rapidly washed in ice cold PBS and flash frozen prior to lysis as described for western blotting. 2 μL lysate was dot blotted onto 0.45 μM nitrocellulose with mouse anti-puromycin (clone 12D10/MABE343; Sigma-Aldrich) antibody at 1:25,000 was used to quantify puromycin incorporation versus a loading control (anti-vinculin, 1:2000; CST), with ImageJ used for quantification.

### Profiling collided ribosomes using sucrose density gradient fractionation

HEK293T cells were plated the day before the experiment at a density of 9 million cells per 15 cm plate. Cells were washed on ice with 20 mL PBS with calcium and magnesium per plate. 500 μL lysis buffer (50 mM HEPES pH 7.4, 100 mM KOAc, 15 mM Mg(OAc)₂, 5% ultrapure glycerol, 0.25% NP-40, 1× Halt protease/phosphatase inhibitor cocktail, 20 U/mL Turbo DNase, 1 mM DTT, 100 µg/mL cycloheximide, 5 mM CaCl₂) was added to plates on ice and lysate was collected. Lysate was transferred to an eppendorf tube, triturated with a 26G needle 20 times, and clarified by spinning at 8000 x *g* for 5 minutes. The supernatant was digested with 4.4 μL MNase (Worthington Biochemical) / 500 μL lysate at 16°C for 1 hour with shaking. Sucrose density gradients (10% or 50% sucrose in 25 mM HEPES pH 7.4, 100 mM KOAC, 5 mM Mg(OAc)_2_, 1 mM DTT, 100 ug/mL cycloheximide) were created using a BioComp Gradient Station in Seton Polyclear tubes (Seton 7030) and then chilled. Gradients were balanced to 0.01 grams by removing sucrose solution from the top and 500 μL lysate was gently layered on top. Gradients were spun in an SW41 rotor in at 35,000 RPM for 2.5 hours, and fractionation was performed using an BioComp gradient fractionator, measuring A260 nm and collecting 14 fractions per gradient.

Protein was TCA precipitated from sucrose density gradient fractions by adding ∼12% TCA, inverting briefly to mix and incubating at -20°C overnight. Protein was pelleted by spinning at 16,000 x *g* for 30 minutes at 4°C. Supernatant was removed by pipetting, and samples were washed 3x with ice cold acetone, spinning for 10 minutes at 4°C in between each wash. Samples were air dried at room temperature for 15 minutes. 5 uL 10 mM Tris HCl pH 8 was added to rehydrate samples before resuspending in 100 μL 1x LDS sample buffer with 5 mM DTT. Before western blotting, samples were boiled for 15 minutes at 95°C. 20 μL was loaded per fraction for western blot.

### Cell viability assays

HEK293T and HCT116 cultured in complete DMEM were collected after trypsinization by centrifugation at 300*g* for 5 min, washed 2× in PBS (Corning 21–040 CV) and resuspended in PBS at 1.5 million/mL. Cells were then added to amino acid limitation medium at a final concentration of 9,375 cells per mL. Forty μL of cell suspension was pipetted into the wells of 384-well microplates (Thermo Fisher Scientific 164610). Each data point was present in 10 replicates, and five separate identical plates were prepared for viability measurement after 0, 3, 6, 9 and 12 days incubation. Assay plates were incubated at 37°C/5% CO2 inside containers humidified by sterile wet gauze. Plates were removed from incubation and cooled at room temperature for 10 min, before dispensing 40 μl of CellTiter-Glo (CTG) (Promega) into each well. Following a 10 min incubation at room temperature, luminescence was measured in a plate reader (BioTek Synergy H1) (see Fig. S1B).

### Cell proliferation

To quantify doublings per day, 50,000 cells were plated in a 6 well plate in 3 replicate wells plus a parallel plate for counting at experiment onset. The next day, cells were washed and medium exchanged for amino acid limitation or treatment medium. The parallel plate was trypsinized in 500 μL, quenched in 500 μL, and 500 μL cell suspension was counted in 9.5 mL IsoTon II Diluent (Beckman Coulter) using a Beckman Coulter Multisizer in Accuvette cups (Beckman Coulter) to quantify cell number at experiment onset. After 4 days of treatment, cells were similarly counted. We calculated the total number of doublings as log₂(final cell count / initial cell count) and divided this value by 4 to obtain the average doubling rate per day.

### 35S chase assay for protein turnover

HEK293T cells were seeded at 500,000 cells per well in 6 well plates. The next day, cells were washed with PBS and medium replaced with arginine limitation or rich medium. After 4 days of arginine limitation or culture in rich conditions, cells were washed twice in warm arginine limitation medium without methionine, and medium was replaced with 1 mL of arginine limitation or rich medium containing 2 μL radiolabeled EasyTag EXPRESS35S Protein Labeling Mix with L-^35^S-Cysteine and L-^35^S-Methionine (∼11 uCi/μL when fresh; Perkin Elmer) per well. After 30 minutes incubation at 37°C, cells were washed twice in warm treatment medium and then incubated in treatment medium for a “chase” period. At designated time points from 0-100 hours, cells were washed in ice cold PBS and flash frozen in liquid nitrogen. Wells were lysed in 60 μL RIPA buffer per well as described in Western blotting. 20 μL clarified lysate was spotted onto Whatman Grade GF/C glass microfiber filters and allowed to dry for 10-15 minutes. Protein was precipitated by submerging in cold 5% TCA, shaking for 5 minutes at 4°C, and resting for 3 minutes at 4°C. Filters were washed twice for 5 minutes each with cold 10% TCA with shaking at 4°C, twice for 2 minutes with cold ethanol with shaking at 4°C, and once for 2 minutes with cold acetone with shaking at 4°C. Filters were then allowed to air dry at room temperature in a chemical hood, and after drying added to scintillation vials with 5 mL scintillation fluid (EcoScint; National Diagnostics) per vial. Vials were shaken to mix and counted using a scintillation counter (Beckman Model LS 6000LL) for 1 minute. Values were normalized to radioactive counts at the onset of the chase period (0 hours) and plotted over time to assess radioactive protein decay kinetics.

### Quantification of intracellular amino acid levels using GC/MS

Cells were seeded at 500,000 cells per well in 6 well plates. The next day, cells were washed with PBS and medium replaced with amino acid limitation medium. Across a series of 8 time points from 15 min – 5 days, cells were rapidly rinsed twice with ice-cold blood bank saline and extracted using 500 μL ice-cold 80% methanol in water with 4 μg/mL norvaline (Sigma-Aldrich N7627) and 6.25 μM ^13^C-labeled amino acid spike in MSKA2-1.2 (Cambridge Isotope labs) per sample. The soluble supernatant after spinning extracts at 21000 x *g* for 10 min at 4°C was dried overnight under nitrogen gas. Polar metabolites were derivatized to form methoxime-tBDMS derivatives by incubation with 24 μL 2% methoxylamine hydrochloride in pyridine (ThermoFisher TS–45950) and heating at 37°C for 1 h, followed by addition of 30 μL N–methyl–N–(tert–butyldimethylsilyl) trifluoroacetamide +1% tert–Butyldimethylchlorosilane (Sigma-Aldrich 375934) and heating at 80°C for 2 h. Derivatized samples were analyzed by GC/MS using a DB-35MS column (30 m × 0.25 mm i.d. × 0.25 μm, Agilent J&W Scientific) installed in an Agilent 7890B gas chromatograph linked to an Agilent 5977B mass spectrometer. Peaks were confirmed using the ^13^C-labeled amino acid standard and metabolite ion counts determined by integrating characteristic ion fragments for each amino acid derivative (NIST Chemistry WebBook) using the El Maven software v11.0 (Elucidata), corrected for natural isotope abundance using the R package IsoCorrectoR, and normalized to 1) the mean-normalized internal norvaline standard ion count and 2) a cell count collected from an identical 6-well plate processed in parallel.

### FACS-based CRISPRi/a screen

Genome-wide CRISPRi and CRISPRa sgRNA libraries (hCRISPRi_v2: Addgene #83969 and #83970; hCRISPRa_v2: #83978 and #83979) were amplified, packaged into lentiviral particles, and titers were determined as described in ref ^118^. For K562, CRISPRi parental cells (187 million) were infected at a multiplicity of infection (MOI) of 0.28 and selected using 2 μg/mL puromycin (Sigma) for 6 days starting 48 h post-transduction. Cells were maintained at ≥ 250× library coverage at all steps. After 24 h in RPMI rich, cells were starved 72 h in RPMI without arginine. 80 million cells were pelleted by centrifugation, resuspended in PBS, and sorted by FACS (BD Aria II) into high (18 million cells) and low (42 million cells) responder bins (higher than average vs lower than average YFP:mCherry ratio, respectively). Gating was determined using positive and negative reporter activation control cell lines. Sorted cells were recovered and amplified for 5 days in RPMI rich, then re-starved and sorted. The second sort yielded 41 million low responders from the initial low responder sorted population and 3.3 million high responders from the initial high responder sorted population. Cells were allowed to recover in RPMI for 5-6 days, harvested by centrifugation, washed with PBS and pellets stored at −80 °C.

A similar procedure was followed for 293T cells. CRISPRi and CRISPRa parental cells (300 million) were infected at an MOI of 0.30-0.32 and selected using 2 μg/mL puromycin (Sigma) for 5 days starting 48 h post-transduction. After 24 h in DMEM rich medium, 150 million cells were plated in DMEM without arginine and starved for 72 h. Cells (310-340 million) were then harvested by trypsinization, quenched with DMEM without arginine, pelleted, resuspended in PBS, and sorted into high (40-44 million cells) and low (40-44 million cells) responder bins (top and bottom 20% YFP:mCherry ratio). After 4-5 days recovery, 100 million cells from each bin were re-starved for 72 h, and 27-37 million cells were collected in each condition. Cells were allowed to recover in DMEM for 4 days, washed once with PBS and pellets were stored at −80°C.

Genomic DNA was extracted from sorted cells and sgRNA counts were determined as previously reported^117^. Briefly, sgRNAs were amplified by PCR using Q5 polymerase (NEB), gDNA extracted using QiaAmp DNA Blood Midi kit (Qiagen), and dual indexed/staggered primers. PCR products were purified by agarose gel electrophoresis, pooled and sequenced on an Illumina NextSeq 500 platform using 75 bp single-end reads.

Screen data were processed using a published pipeline^119^. For each sgRNA, enrichment/depletion was calculated as the abundance in the high responder pool divided by abundance in the low responder pool, normalized to the median of non-targeting controls. For each gene (10 sgRNAs), a phenotype score was defined as the mean of the 7 sgRNAs with the strongest absolute enrichment/depletion values. Gene-level *P* values were determined by comparing the 10 sgRNAs targeting each gene with the 3,790 non-targeting control (NTC) sgRNAs using a Mann–Whitney test. sgRNAs with fewer than 25 reads in either bin were excluded. A pseudocount of 10 was added to all sgRNAs to mitigate low-count noise. Screen scores were calculated by multiplying phenotype scores by −log_10_(*P* value). To estimate technical noise, negative control “genes” (the same number as that of real genes) were computed by randomly grouping 10 sgRNAs from the pool of NTC sgRNAs. False discovery rates were then calculated by comparing real and control gene distributions.

### Charged tRNA sequencing

To sequence charged tRNA, we utilized the method described in ref ^120^. 1.8 million HEK293T cells were plated in a 10 cm plate. The next day, cells were washed with PBS and treated with 15 mL amino acid limitation medium for the designated times between 15 minutes and 5 days (see Figure 2). Plates were then washed with PBS and flash frozen in liquid nitrogen. RNA isolation was performed with TRIzol and isopropanol precipitation. 5-10 μg total RNA in 100 mM sodium acetate pH 4.5, 1 mM EDTA, was treated with spike in E. coli tRNA controls, oxidized with sodium metaperiodate, subject to beta hydrolysis using lysine pH 8, dephosphorylated using rSAP (NEB), and cleaned up using the Monarch RNA cleanup kit (NEB). tRNA was size selected on a 10% TBE Urea gel (Novex). 500 ng to 2 μg tRNA was adaptor ligated, reverse transcribed using Maxima reverse transcriptase (Thermo Scientific), circularized using CircLigase (Lucigen), and amplified by PCR with KAPA HiFi polymerase (Roche) for 8-12 cycles to add library barcodes for Illumina paired end sequencing (2x100 bp reads). Data analysis was performed using the python pipeline published on Github^120^ (https://github.com/krdav/tRNA-charge-seq/tree/main). Output csv files were then processed using R in jupyter notebook to filter tRNAs out with less than 100 reads, calculate absolute abundance of charged tRNA (reads per million [RPM] * fraction CCA-ending reads), sum or average across all isodecoder tRNAs for each isoacceptor family, normalize to rich controls to determine changes upon amino acid limitation, and plot various metrics for cognate tRNAs for each amino acid limitation condition.

### Proteomics

#### Sample collection for steady-state protein abundance measurements

HEK293T cells were cultured for six days in lysine-free DMEM supplemented with 800 µM ^13^C₆-lysine (Cambridge Isotope Laboratories, CLM-2247-H-0.1) to generate “heavy” samples, or in standard DMEM for the same duration to generate “light” samples. After labeling, approximately 7 × 10^5^ cells were seeded per 6 cm dish in the corresponding labeling medium and allowed to adhere overnight. Cells were then washed once with phosphate-buffered saline (PBS) and incubated in either amino acid limitation medium (see *Preparation of amino acid dropout medium* section) or nutrient-rich control medium. After four days of treatment, cells were trypsinized, counted, and harvested by centrifugation (200 × g, 5 min). For proteomic analysis, approximately 2 × 10^^5^ amino acid–limited cells and 1 × 10^6^ nutrient-rich control cells were washed once in PBS and flash-frozen as pellets in liquid nitrogen. Amino acid-limited samples were mixed with nutrient-rich control samples from the opposite isotopic channel. A replicate assay was performed with isotopic labels swapped between the amino acid-limited and rich medium conditions to control for isotope-label-specific effects.

In parallel, an additional pellet of identically treated cells was collected for RNA sequencing. Total RNA was isolated using the Zymo RNA Miniprep kit and processed for Illumina-based library preparation and sequencing to enable calculation of protein abundance per mRNA.

#### Sample collection for pulsed SILAC to measure protein synthesis rates

For pulsed SILAC experiments, approximately 2 × 10^6^ HEK293T cells were seeded in 10 cm dishes for amino acid limitation conditions, and approximately 5 × 10^5^ cells were seeded for nutrient-rich control conditions. The following day, cells were washed once with PBS and incubated in 15 mL of either amino acid limitation medium or nutrient-rich medium per dish.

After two days under nutrient-rich conditions or four days under amino acid limitation, cells were washed with PBS and the medium was exchanged for the same condition in which lysine was fully replaced with ^13^C₆-lysine. Cells were harvested at 0, 4, 8, 24, and 48 hours following isotope addition. At each time point, cells were trypsinized, counted, and two-thirds of the population was pelleted, washed once with PBS, and flash-frozen in liquid nitrogen for proteomic analysis.

As an internal quantitation reference, HEK293T cells fully labeled for at least seven doublings in medium bearing ^13^C₆^15^N₂-lysine (Cambridge Isotope Labs CNLM-291-H-0.5); these cells were then prepared separately and added as a spike-in control to each sample to facilitate quantitative comparison across samples and biological replicates, following ref ^121^.

#### Sample processing for LC-MS/MS analysis

Frozen cell pellets were resuspended in water and lysed in buffer containing 5% (w/v) sodium dodecyl sulfate and 50 mM triethylammonium bicarbonate, pH 8.5 (TEAB; Sigma Aldrich #18597). Proteins were reduced with 10 mM 1,4-dithiothreitol (CAS: 3483-12-3) for 30 min at room temperature and alkylated with 10 mM iodoacetamide for 10 min in the dark. Tryptic peptides were generated and purified S-Trap™ micro columns according to the manufacturer’s protocol, with minor modifications as follow. Proteins were digested on-column using sequence-grade trypsin and Lys-C reconstituted in 50 mM TEAB, with the protease added to the column at an enzyme-to-protein ratio of 1:10 (w/w). The digestion was carried out for 2 hours at 42 °C. Following digestion, peptides were eluted, dried in a speed-vac, and resuspended in MS sample buffer consisting of 4% acetonitrile and 0.1% formic acid.

### LC-MS/MS data acquisition

Peptides (∼2 µg per injection) were analyzed using an Ultimate 3000 UHPLC system coupled to either a Q-Exactive HF or Orbitrap Exploris 480 mass spectrometer (Thermo Fisher Scientific). Samples were loaded onto a PepMap100 C18 precolumn and separated on a PepMap C18 analytical column (75 µm × 25 cm, 3 µm particles, 100 Å pore size) using a 135 min linear gradient from 4% to 30% acetonitrile in 0.1% formic acid at a flow rate of 300 nL min⁻¹.

For data-dependent acquisition, MS1 scans were acquired over an m/z range of 350–1400 at 60,000 resolution (AGC target 3 × 10^6^, maximum injection time 50 ms). The top 12 precursor ions were fragmented by higher-energy collisional dissociation (HCD) at 25% normalized collision energy, and MS2 spectra were acquired at 15,000 resolution (AGC target 1 × 10^5^, maximum injection time 100 ms, isolation window 2.2 m/z).

For data-independent acquisition, MS1 scans were acquired at 120,000 resolution, followed by MS2 scans acquired at 30,000 resolution across 25 variable-width isolation windows^122^ using HCD at 25% normalized collision energy.

### Initial data processing and normalization

DIA datasets were analyzed using Spectronaut Pulsar (v15–16; Biognosys) with the DirectDIA workflow against the human protein database (UP05640). DDA datasets were included as a hybrid spectral library. Searches included three lysine label states (unlabeled, Lys6, Lys8), carbamidomethylation of cysteine as a fixed modification, and oxidation of methionine as a variable modification. In silico generation of missing label channels was enabled.

Precursor intensities were quantified at the MS1 level by integration of extracted ion chromatograms. Heavy-labeled precursors were quantified using M0 and M+1 isotopomers, whereas spike-in control precursors were quantified using M+1 and M+2 isotopomers. All precursor intensities were normalized to the internal spike-in control according to:

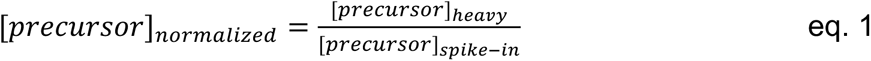

Only peptides with C-terminal lysine residues were considered. Protein abundance was calculated as the median of corresponding peptide values, and proteins quantified by fewer than two peptides were excluded from analysis.

### Kinetic modeling of protein synthesis

Protein synthesis rates were determined by fitting normalized precursor intensities over time to an integrated rate equation (eq. 2) describing the accumulation of heavy isotope–labeled protein as a function of synthesis and degradation rates:

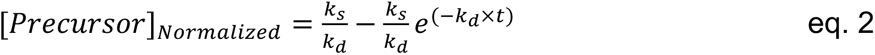

where *k_s_* represents the protein synthesis rate constant, *k_d_* represents the protein degradation rate constant, and *t* denotes time.

Model fits were evaluated using the coefficient of determination (eq. 3).

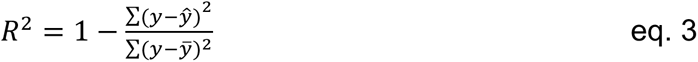

where, *y* is the observed normalized precursor intensity, *ŷ* is the fitted value and *ȳ* is the mean of the observed data. Only proteins with fitted models exhibiting *R^2^* > 0.7 were retained for downstream analysis.

### Proteomics data analysis

The change in steady state protein abundance per mRNA upon amino acid limitation was calculated by subtracting the log_2_ fold change in protein abundance upon amino acid limitation from the log_2_ fold change in mRNA level (DESeq2) upon amino acid limitation. This was plotted against arginine codon frequency per mRNA (AGA/AGG/CGA/CGC/CGG/CGU codons per total codons) and a Spearman correlation coefficient and p-value were calculated using stat_cor() in R.

The log_2_ fold change in protein synthesis rate per mRNA upon amino acid limitation was calculated by subtracting the log_2_ rate in amino acid limited conditions from the log_2_ rate in nutrient rich conditions. This was plotted against arginine codon frequency per mRNA (AGA/AGG/CGA/CGC/CGG/CGU codons per total codons) and a Spearman correlation coefficient and p-value were calculated using stat_cor() in R.

Peptide distribution skew from N to C terminus was calculated as follows. First, we found normalized peptide positions from N to C terminus by taking the start position of each peptide and normalizing it to a fraction of the total length of the protein from 0 to 1; these numbers were rounded to bin peptides into one-tenth increments along the length of a protein. Then at all peptides in each bin, the average log_2_ fold change in protein synthesis rate upon amino acid limitation was calculated. Finally, average log_2_ fold change in protein synthesis rate upon amino acid limitation values were normalized to bin 0, the N terminus, to isolate changes across the length of a protein. These normalized average log_2_ fold change in protein synthesis rate values were plotted against binned peptide position to reveal trends and a Spearman correlation coefficient and p-value were calculated using stat_cor() in R.

To identify transcripts whose translation was likely to be negatively regulated by AGA content upon arginine limitation, we intersected the transcripts in the top 25% of AGA frequency with a list of corresponding proteins whose synthesis rate we could measure, and was reduced upon arginine limitation, in both WT and GCN2 KO HEK293T cells. The resulting list of 118 transcripts was analyzed for GO term enrichment using Enrichr^123^.

### Data analysis and visualization

See individual sections for a description of statistics employed. All data analysis and visualization was performed in R using jupyter notebook and ggplot.

## Acknowledgements

We thank current and former members of the Vander Heiden and Darnell labs for insight and feedback, including M. Luck, B.A. Michel, and C. Barrington-Ham. We thank S. Chang, M. Munim, and M. Bartusel for assisting with radioactivity work, and C. Waldie and R. Sangwan for technical assistance. We thank F. Diehl and S. Kim for sharing protocols and reagents. We thank S. Kim, A. Subramaniam, J. Weissman, R. Green., J. Rutter, R. Darnell, A. Muir, E. Lien, Z. Li, and F. Diehl for helpful discussions throughout the project and for input on the manuscript. We thank G. Paradis and the Koch Institute Flow Cytometry Core for training, assistance, and ensuring a consistently high-quality standard for flow analysis and sorting. We thank the MIT BioMicro Center for assistance with RNA library prep and sequencing, and the Fred Hutch Genomics and Bioinformatics Facility for assistance with sequencing of charged tRNA seq libraries.

This work was supported in part by the Koch Institute Cancer Center Support Grant P30CA14051. A.M.D was supported by a Jane Coffin Childs Memorial Foundation Postdoctoral fellowship and a Charles A King Postdoctoral Fellowship. C.C. and P.K.S were supported by the Harvard Ludwig Cancer Center. L.B.S was supported by the NIH (R35GM147118 and P30CA015704). J.H.D acknowledges support from the NIH (R00AG050749) and the Smith Family Foundation Odyssey Award. M.G.V.H. acknowledges support from the MIT Center for Precision Cancer Medicine, the Ludwig Center at MIT, and the NIH (R35CA242379).

## Data Availability

All sequencing data will be deposited on GEO (pending). All proteomics data is deposited on MASSive (username: MSV000099718_reviewer; password: rqc). All processed charged tRNA seq data, proteomics, and genome-wide CRISPR screen data is compiled in the following Github repository (https://github.com/adarnell/2026_Darnell_translationDisruptionManuscript/). Full western blots and flow cytometry time courses for translation disruption reporter survey are provided as supplementary files (S1-S5).

## Contributions

Conceptualization: A.M.D., C.C., M.G.V.H.; Methodology: A.M.D., C.C., V.P., K.D., D.S.C., J.H.D., M.G.V.H.; Investigation: A.M.D., C.C., V.P., K.D., D.S.C., K.L.A., R.E., C.P.V.H.; Formal analysis: A.M.D, C.C, K.D., D.S.C.; Resources: L.B.S., P.K.S., J.H.D.; Writing – Original Draft: A.M.D.; Writing – Review & Editing: All authors; Supervision: M.G.V.H., J.H.D., P.K.S.; Funding acquisition: M.G.V.H., P.K.S., J.H.D.

## SUPPLEMENTARY FIGURE LEGENDS

**Figure S1:**
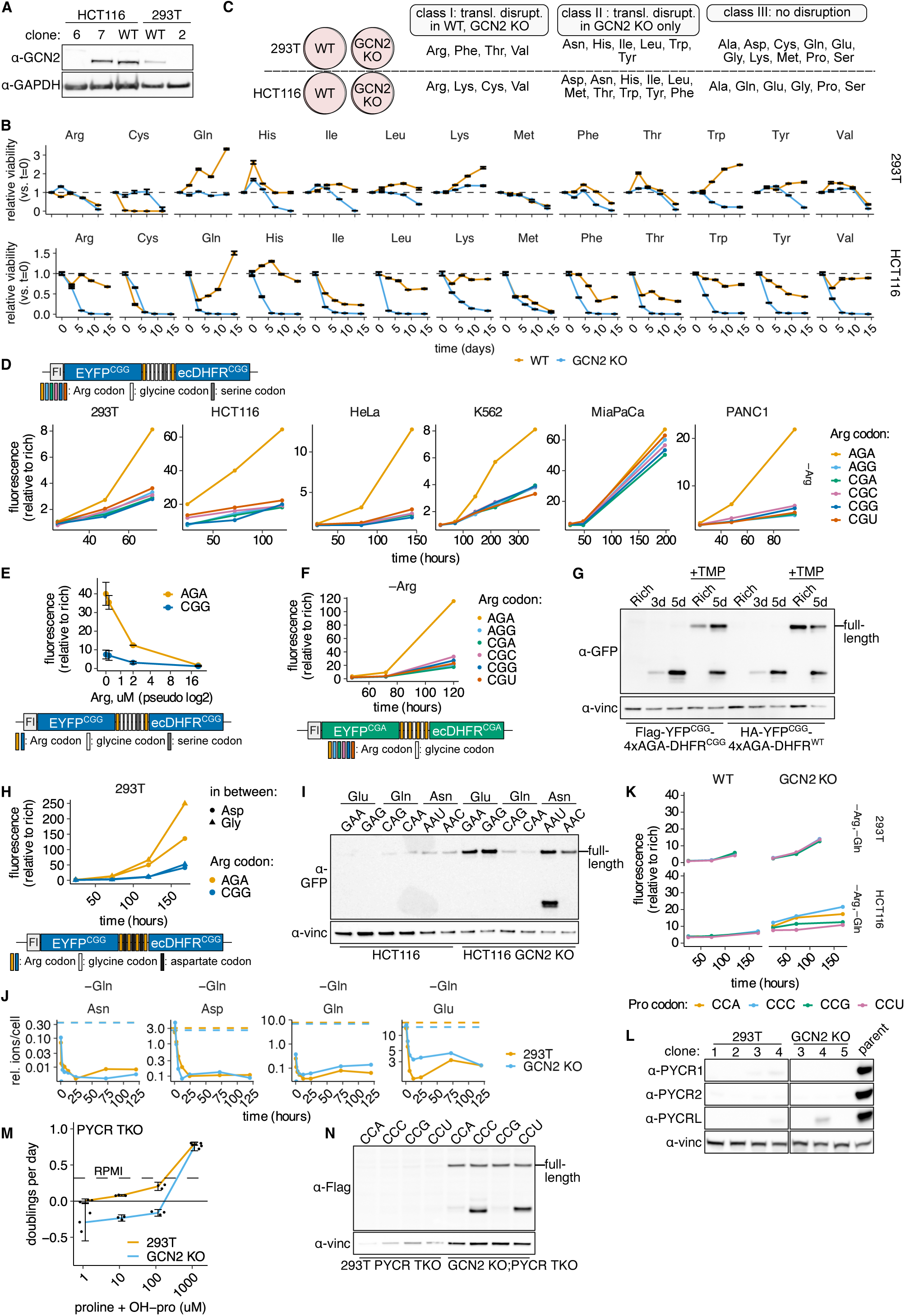
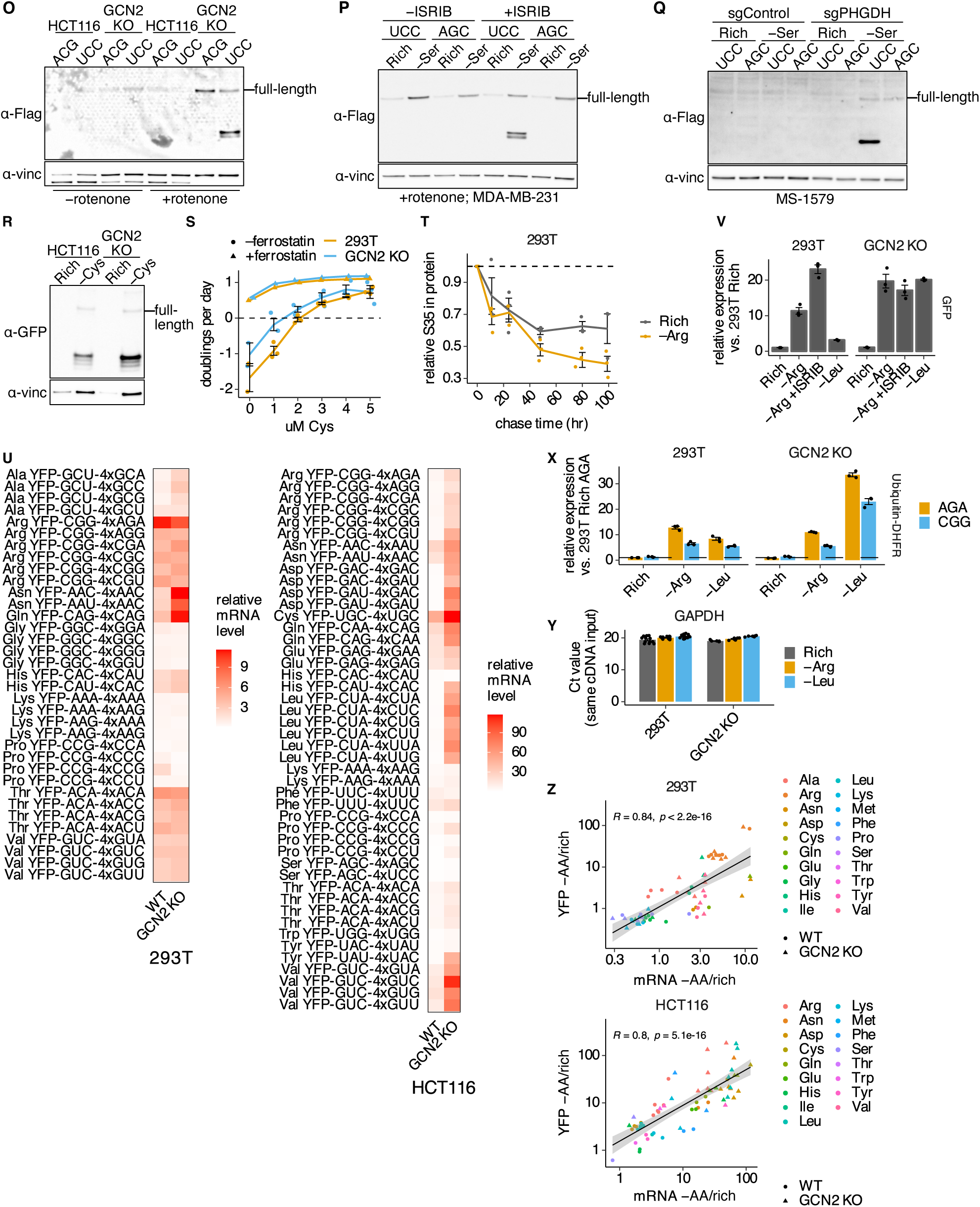
A survey of translation disruption across codons and conditions reveals that arginine AGA codons are a unique trigger. (A) Western blot to confirm HCT116 and HEK293T (293T) GCN2 KO clones. Lane labels correspond to clone numbers. Subsequent experiments used HCT116 GCN2 KO clone 6 and 293T GCN2 KO clone 2. (B) Cell viability changes over time upon limitation for the indicated amino acid in wildtype (WT) and GCN2 KO 293T or HCT116 cell lines as indicated. Dashed line indicates value for no change in cell number. (C) Summary of survey results; the survey was performed in WT and GCN2 KO 293T or HCT116 cells and amino acids (AAs) were categorized into class I/II/III as described based on observed translation disruption (transl. disrupt.) patterns. (D) Modified arginine codon translation disruption reporter (Flag-YFP^CGG^-2xAGA-DHFR^CGG^) change in fluorescence upon arginine limitation as measured by flow cytometry across 6 cell lines. (E) Modified arginine codon translation disruption reporter (Flag-YFP^CGG^-2xAGA-DHFR^CGG^ or Flag-YFP^CGG^-2xCGG-DHFR^CGG^) change in fluorescence after 6 days of culture at the indicated arginine concentration measured by flow cytometry. (F) Arginine codon translation disruption reporter (with CGA codons in YFP/DHFR; Flag-YFP^CGA^-4xAGA-DHFR^CGA^) change in fluorescence upon arginine limitation as measured by flow cytometry. (G) Western blot to detect translation disruption product accumulation upon limitation for arginine for 3 or 5 days (d) or in rich conditions, +/−TMP, in cells expressing either Flag-YFP^CGG^-4xAGA-DHFR^CGG^ or HA-YFP^CGG^-4xAGA-DHFR^WT^ as indicated. (H) Change in fluorescence of the arginine codon translation disruption (Flag-YFP^CGG^-4xAGA-DHFR^CGG^) or control (Flag-YFP^CGG^-4xCGG-DHFR^CGG^) linker variants with either GGA glycine codons or GAC aspartate codons in between the linker Arg codons upon arginine limitation as measured by flow cytometry. (I) Western blot to detect translation disruption product accumulation for glutamine (Flag-YFP^CAG^-4xCAA-DHFR or Flag-YFP^CAA^-4xCAG-DHFR), asparagine (Flag-YFP^AAC^-4xAAU-DHFR or Flag-YFP^AAU^-4xAAC-DHFR), and glutamate reporters (Flag-YFP^GAG^-4xGAA-DHFR or Flag-YFP^GAG^-4xGAG-DHFR) in wild-type and GCN2 KO HCT116 cells upon limitation for glutamine for 4 days as indicated; lane label indicates linker codon identity. (J) GC/MS analysis of relative intracellular asparagine, aspartate, glutamine, glutamate levels over time following glutamine limitation in wild-type and GCN2 KO 293T cells. Dashed line indicates intracellular amino acid level in rich medium for each cell line. Ion counts are internally normalized to a norvaline spike-in and cell count. (K) Flow cytometry to detect reporter fluorescence in wild-type and GCN2 KO 293T cells or HCT116 cells expressing proline translation disruption reporters (Flag-YFP^CCG^-4xCCA/C/G/U-DHFR) over time upon limitation for proline, arginine, and glutamine. (L) Western blots for PYCR1, PYCR2, and PYCRL (PYCR3) to identify 293T and GCN2 KO triple KO clones (PYCR TKO). Subsequent experiments use 293T PYCR TKO clone 1 and GCN2 KO PYCR TKO clone 3. (M) Doublings per day of selected PYCR TKO clones in wild-type and GCN2 KO 293T cells across a titration of proline and 3-hydroxy-L-proline levels. (N) Western blot to detect translation disruption product accumulation in control (293T) and GCN KO PYCR TKO 293T cells expressing proline translation disruption reporters (Flag-YFP^CCG^-4xCCA/C/G/U-DHFR) upon limitation for proline, for 4 days. (O-Q) Western blots to detect translation disruption product accumulation in HCT116 (O), MD-MBA-231 (P), or MS-1579 (Q) cells expressing serine translation disruption (Flag-YFP^AGC^-4xUCC-DHFR) and control (Flag-YFP^AGC^-4xAGC-DHFR) reporters^5^ upon limitation for serine and glycine with or without 80 nM rotenone for 2 days (O), limitation for serine with or without 40 nM ISRIB for 5 days (P), or limitation for serine for 3 days (Q). (R) Western blot to detect translation disruption product accumulation in HCT116 cells expressing cysteine translation disruption reporter (Flag-YFP^UGC^-4xUGU-DHFR) upon limitation for cysteine for 5 days. (S) Doublings per day of wild-type and GCN2 KO 293T cells across a titration of cysteine levels, with or without 2 μM ferrostatin. (T) Decay of ^35^S-Met incorporated into total cellular protein in arginine limited or rich conditions over time. Dashed line represents normalized ^35^S signal at chase start. (U) mRNA level of indicated translation disruption reporter (“YFP-NNN” refers to codons used to encode YFP, followed by 4xNNN linker) measured by RT-qPCR after limitation for the cognate amino acid for 2-7 days (corresponding to the time point of maximum flow cytometry reporter signal) relative to mRNA level in rich conditions in wild-type (WT) and GCN2 KO 293T or HCT116 cells. (V-X) mRNA levels of EGFP (“GFP”) (V) or HA-ubiquitin-DHFR (X) transgenes measured by RT-qPCR in wild-type and GCN2 KO 293T cells upon limitation for leucine or arginine and/or treatment with 40 nM ISRIB for 5 days, normalized to 293T Rich sample expressing the Flag-YFP^CGG^-4xAGA-DHFR^CGG^ (“AGA”) or Flag-YFP^CGG^-4xAGA-DHFR^CGG^ (“CGG”) reporter as indicated. (Y) C_t_ value of endogenous GAPDH mRNA given equal cDNA inputs for RT-qPCR upon arginine or leucine limitation for 5 days or rich conditions in wild-type and GCN2 KO 293T cells. (Z) Correlation between YFP reporter signal and corresponding mRNA level change upon limitation for the cognate amino acid (−AA/rich), log_10_ scaled axes. Spearman correlation coefficients (R) and associated p-values are shown on the plot. (G,I,L,N-R) Full-length reporter product is indicated (α-GFP antibody used to detect YFP, α-vinc = α-vinculin as a loading control).

**Figure S2:**
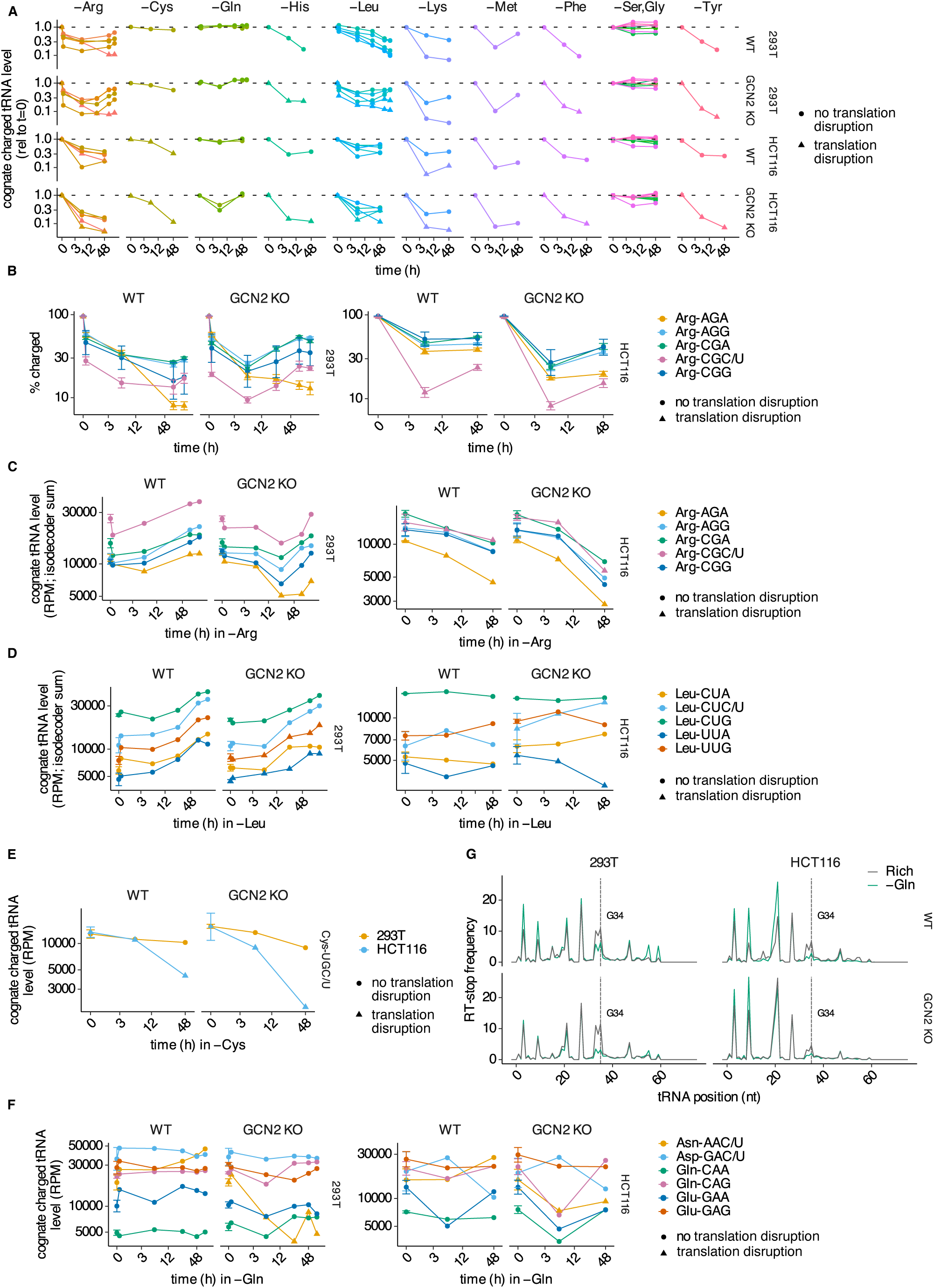
Isoacceptor-specific charged tRNA depletion underlies codon-specific translation disruption. (A) Charged tRNA levels for the codons cognate to the limiting amino acid(s) over time in wild-type (WT) and GCN2 KO 293T or HCT116 cells as indicated, normalized to time = 0. (B-E) Percent tRNA charged (B), total tRNA levels (reads per million), or charged tRNA levels (reads per million) for the arginine (B,C), leucine (D), cysteine (E), or glutamine, glutamate, aspartate, and asparagine (F) isoacceptor tRNAs cognate to the indicated codons upon limitation for the corresponding amino acid in WT and GCN2 KO 293T or HCT116 cells as indicated. Error bars in C-F are only present at t=0 and reflect standard error of the mean from n=3 biological replicate measurements, error bars in B represent the reflect standard error of the mean over all isodecoder tRNAs. (G) Reverse transcription (RT) stop frequency as a proxy for the frequency of specific modifications at each nucleotide (nt) in the Asp-tRNA^GUU^ tRNA in WT and GCN2 KO 293T or HCT116 cell lines in rich conditions or upon glutamine limitation for 5 days in 293T cells or 2 days in HCT116 cells; dashed vertical line marks predicted queuosine-modified G34 position. (B-D,F) For tRNAs that decode more than one codon through wobble base pairing, multiple codons are listed.

**Figure S3:**
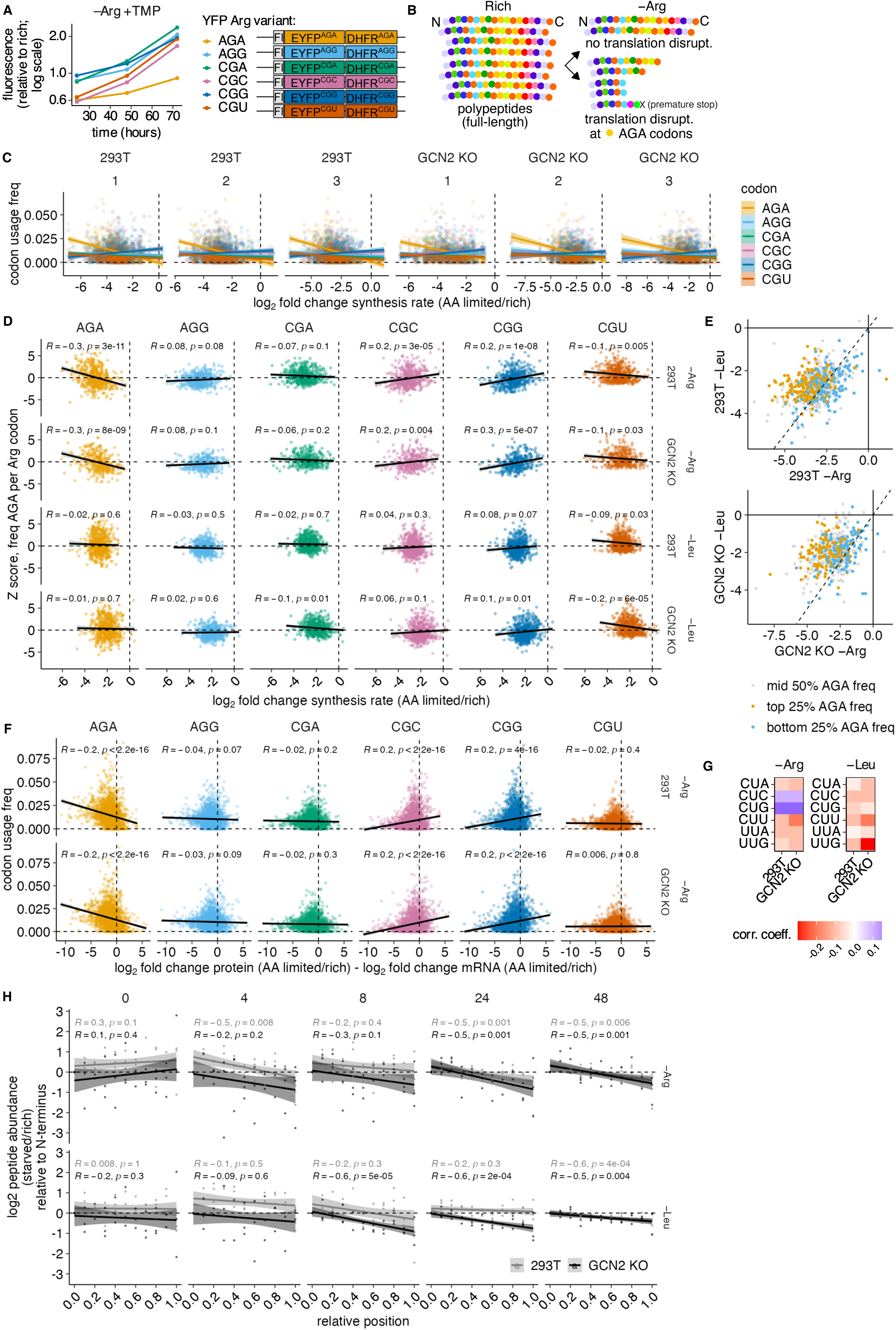
Translation disruption at AGA codons reduces endogenous protein production. (A) YFP-DHFR codon variant TMP-stabilizable protein synthesis rate reporter (Flag-YFP-DHFR) fluorescence accumulation upon arginine limitation with concurrent TMP addition in 293T cells, measured by flow cytometry. (B) Schematic of the predicted effect of translation disruption on protein production and peptide distribution as a function of position from N to C terminus. If AGA codons (yellow) reduce protein synthesis rate without causing translation disruption, peptide distribution remains even across polypeptide N to C terminus. If translation disruption occurs, peptides are expected to be depleted progressively towards the C terminus through premature ribosome fall-off or frameshifting. (C-D) Transcript codon usage frequency (C) or Z-score estimating codon usage bias (D) for each arginine codon plotted against the log_2_ fold-change (l2fc) in protein synthesis rate for the corresponding protein upon limitation for arginine or leucine for 4 days in wild-type and GCN2 KO 293T cells as indicated. Regression line, Spearman correlation coefficient (R) and associated p-value are shown on each plot. Note that 5 outlier points with log_2_ fold-change below −7.5 are not shown. In (C), 3 biological replicate measurements are plotted separately. (E) Log_2_ fold-change (l2fc) in protein synthesis rate for the corresponding protein upon limitation for arginine versus leucine for 4 days in wild-type and GCN2 KO 293T cells as indicated. Corresponding transcripts are binned by AGA usage frequency (high = top 25%, mid = 25-75%, low = bottom 25%). Note that 1 outlier point with l2fc below −10 is not shown in the GCN2 KO plot. Dashed line represents y=x. (F) Codon usage frequency for each arginine codon plotted against the log_2_ fold-change (l2fc) in protein abundance per mRNA upon limitation for arginine or leucine for 4 days in wild-type and GCN2 KO 293T cells as indicated. Regression line, Spearman correlation coefficient (R) and associated p-value are shown on each plot. Note that 6 outlier points with l2fc below −10 are not shown. (G) A heatmap of associated Spearman correlation coefficients from comparing transcript codon usage frequency for each leucine codon to the log_2_ fold-change in protein synthesis rate for the corresponding protein upon limitation for arginine or leucine for 4 days in wild-type and GCN2 KO 293T cells as indicated. (H) Log_2_ fold-change (l2fc) in average peptide abundance upon limitation for arginine or leucine plotted against the binned fractional position of the peptide from 0 (N-terminus) to 1 (C-terminus), normalized to abundance at the N-terminus, for all peptides in wild-type and GCN2 KO 293T cells, at each SILAC label incorporation time point (0/4/8/24/48 hours; see Fig. 3A and Methods). In all panels except A-C/F, points are averages across 3 biological replicates.

**Figure S4:**
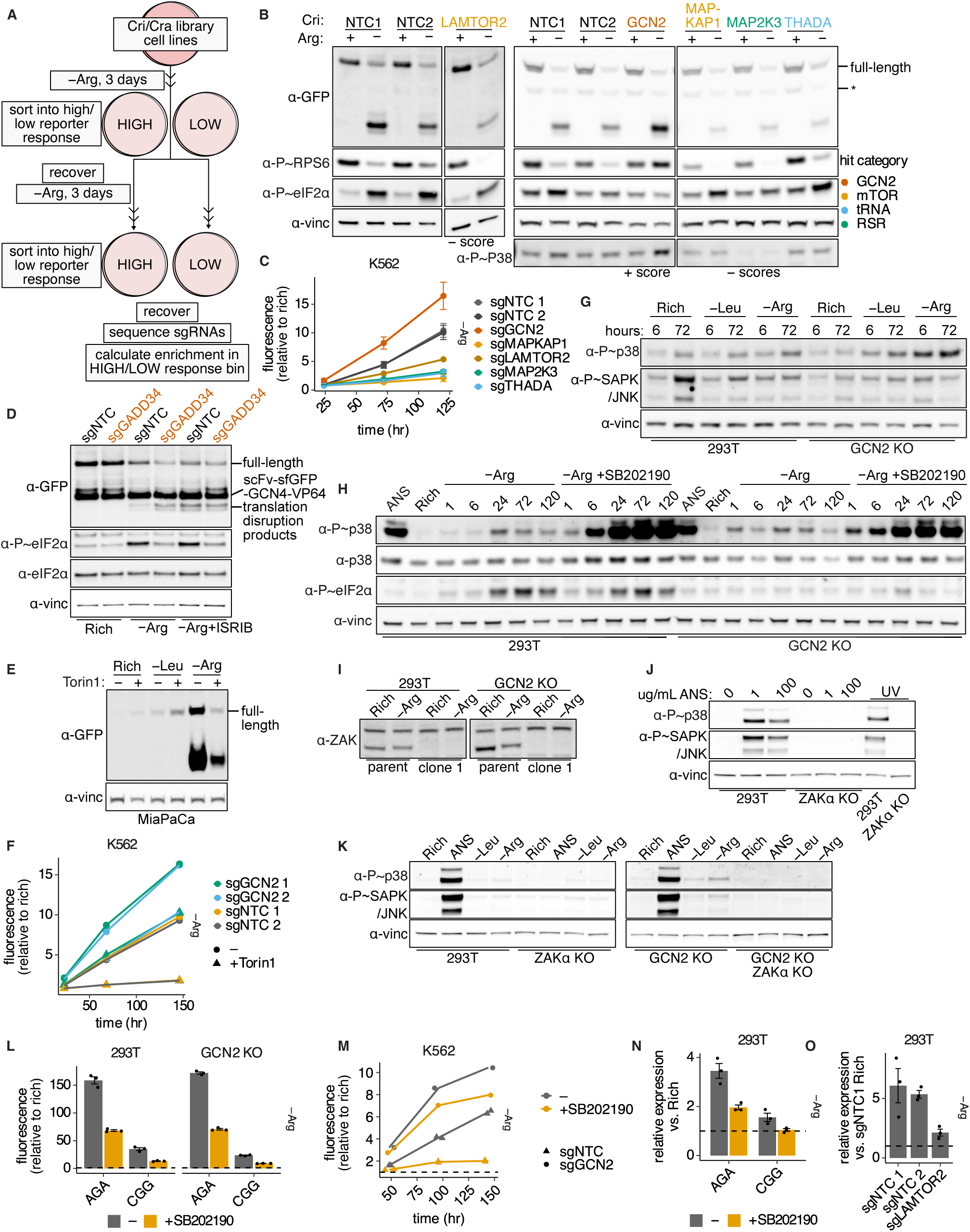
A genome-wide screen reveals the core regulatory network controlling translation disruption upon arginine limitation. (A) Schematic outlining how the screen with reporters and sorting scheme depicted in Fig. 4A was performed. Cells were arginine limited for 3 days, sorted, recovered, and re-sorted into the same bin after a second period of arginine limitation. After recovery, guide RNAs were sequenced to calculate enrichment scores. (B,C) Western blot (B) and flow cytometry (C) validation of selected hits with negative or positive phenotype scores across various pathways in K562 cells. Cells expressing CRISPRi targeting hits (or non-targeting controls; NTC) and dual color translation disruption reporters (Flag-YFP^CGG^-4xAGA-DHFR^CGG^ and Flag-mCherry^CGG^-4xCGG-DHFR^CGG^) were arginine limited for 3 days to assess translation disruption product levels (B,C) and signaling responses through mTOR, GCN2 and ZAKα (B). In (B), “*” marks non-specific band from blot stripping and reprobing. (D) Western blot to assess translation disruption and GCN2 response in 293T cells overexpressing GADD34 or an NTC by CRISPRa (scFv-sfGFP-GCN4-VP64) and dual color translation disruption reporters (Flag-YFP^CGG^-4xAGA-DHFR^CGG^ and Flag-mCherry^CGG^-4xCGG-DHFR^CGG^), with or without limitation for arginine for 5 days and treatment with 40 nM ISRIB. (E) Western blot to assess translation disruption with or without limitation for leucine or arginine for 7 days and treatment with 250 nM Torin1 in MiaPaCa cells expressing the Flag-YFP^CGG^-2xAGA-DHFR^CGG^ reporter. (F) Flow cytometry to assess translation disruption product accumulation upon limitation for arginine with or without GCN2 knockdown by CRISPRi and 250 nM Torin1 treatment in K562 cells expressing the dual color translation disruption reporters (Flag-YFP^CGG^-4xAGA-DHFR^CGG^ and Flag-mCherry^CGG^-4xCGG-DHFR^CGG^). (G-H) Western blots to assess phospho-p38 and -JNK response to arginine or leucine limitation and 0.1 μg/mL anisomycin treatment with or without treatment with 5 μM SB202190 (p38 inhibitor) in wild-type and GCN2 KO 293T cells. (I) Western blot to confirm ZAKα KO in wildtype and GCN2 KO 293T cells as indicated. (J-K) Western blots to assess phospho-p38 and -JNK response to 0.1 μg/mL anisomycin treatment or UV irradiation (J) or arginine or leucine limitation (K) in wild-type and GCN2 KO, ZAKα KO, and ZAKα+GCN2 double KO 293T cells as indicated. (L) Flow cytometry to assess reporter fluorescence upon limitation for arginine for 6 days with or without treatment with 5 μM SB202190 in wild-type and GCN2 KO 293T cells expressing the Flag-YFP^CGG^-4xAGA-DHFR^CGG^ (“AGA”) or Flag-YFP^CGG^-4xCGG-DHFR^CGG^ (“CGG”) reporter as indicated. (M) Flow cytometry to assess reporter fluorescence over time upon limitation for arginine with or without treatment with 5 μM SB202190 in K562 cells with (sgGCN2) or without (sgNTC) GCN2 CRISPRi knockdown expressing the dual color translation disruption reporters (Flag-YFP^CGG^-4xAGA-DHFR^CGG^ and Flag-mCherry^CGG^-4xCGG-DHFR^CGG^). (N) Change in translation disruption reporter (Flag-YFP^CGG^-4xAGA-DHFR^CGG^ (“AGA”) or Flag-YFP^CGG^-4xAGA-DHFR^CGG^ (“CGG”)) mRNA level upon arginine limitation for 3 days with or without treatment with 5 μM SB202190 in 293T cells. (O) Change in translation disruption reporter mRNA level (Flag-YFP^CGG^-4xAGA-DHFR^CGG^) upon arginine limitation for 3 days with (sgLAMTOR2) or without (sgNTC) CRISPRi knockdown of LAMTOR2 in 293T cells expressing the dual color translation disruption reporters (Flag-YFP^CGG^-4xAGA-DHFR^CGG^ and Flag-mCherry^CGG^-4xCGG-DHFR^CGG^), relative to cells expressing control guide (sgNTC 1). (C,L,N,O) Error bars represent standard error of the mean of 3 replicates. (B,D,E, G-K) Full-length reporter product is indicated (α-GFP antibody used to detect YFP, α-vinc = α-vinculin).

**Figure S5:**
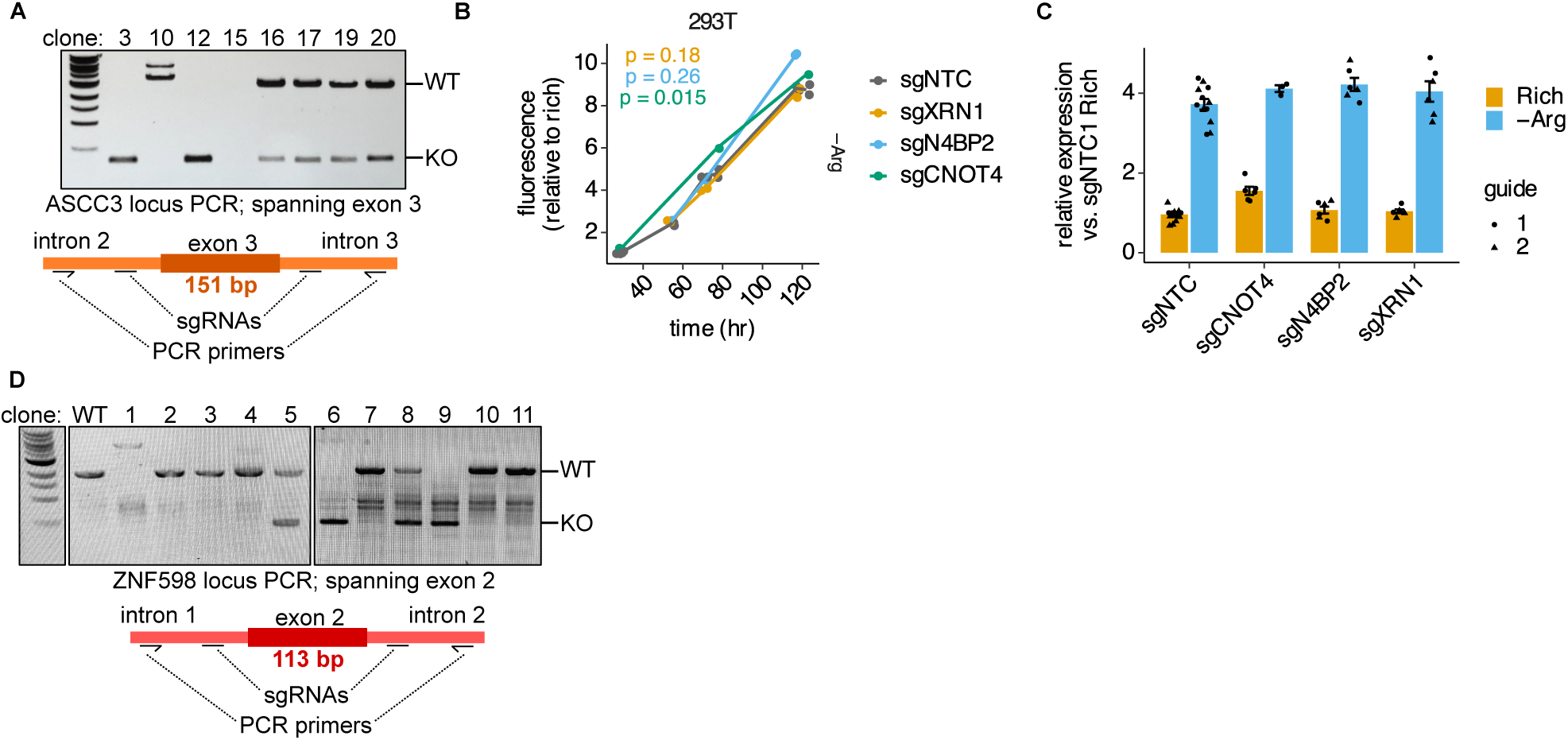
The ribosome splitting and quality control pathways resolve stalled ribosomes during arginine limitation. (A) Validation of ASCC3 KO cell line generation by CRISPR/Cas9. PCR was used to assess ASCC3 locus integrity; successful targeting with 2 sgRNAs leads to exon 3 loss and introduces a frameshift. Clone 3 was used in subsequent experiments. (B,C) Flow cytometry to assess translation disruption product accumulation (B) or RT-qPCR to assess reporter mRNA level (C) in control cells (sgNTC) or upon CRISPRi knockdown of CNOT4, N4BP2, or XRN1 upon limitation for arginine over time (B) or for 3 days (C) in 293T cells expressing the dual color translation disruption reporter integrated at the AASV1 locus. In B) ANOVA was used to determine significance with pairwise differences assessed using estimated marginal means with a Tukey correction for multiple testing where applicable; p-values are shown on the plot. P-values represent sgNTC vs sgGuide comparison and are colored according to guide. In C, error bars represent standard error of the mean across 3-6 biological replicates and two guides. (D) Validation of ASCC3 KO cell line generation by CRISPR/Cas9. PCR was used to assess ZNF598 locus integrity; successful targeting with 2 sgRNAs leads to exon 2 loss and introduces a frameshift. Clones 6 and 9 were used in subsequent experiments.

**Figure S6:**
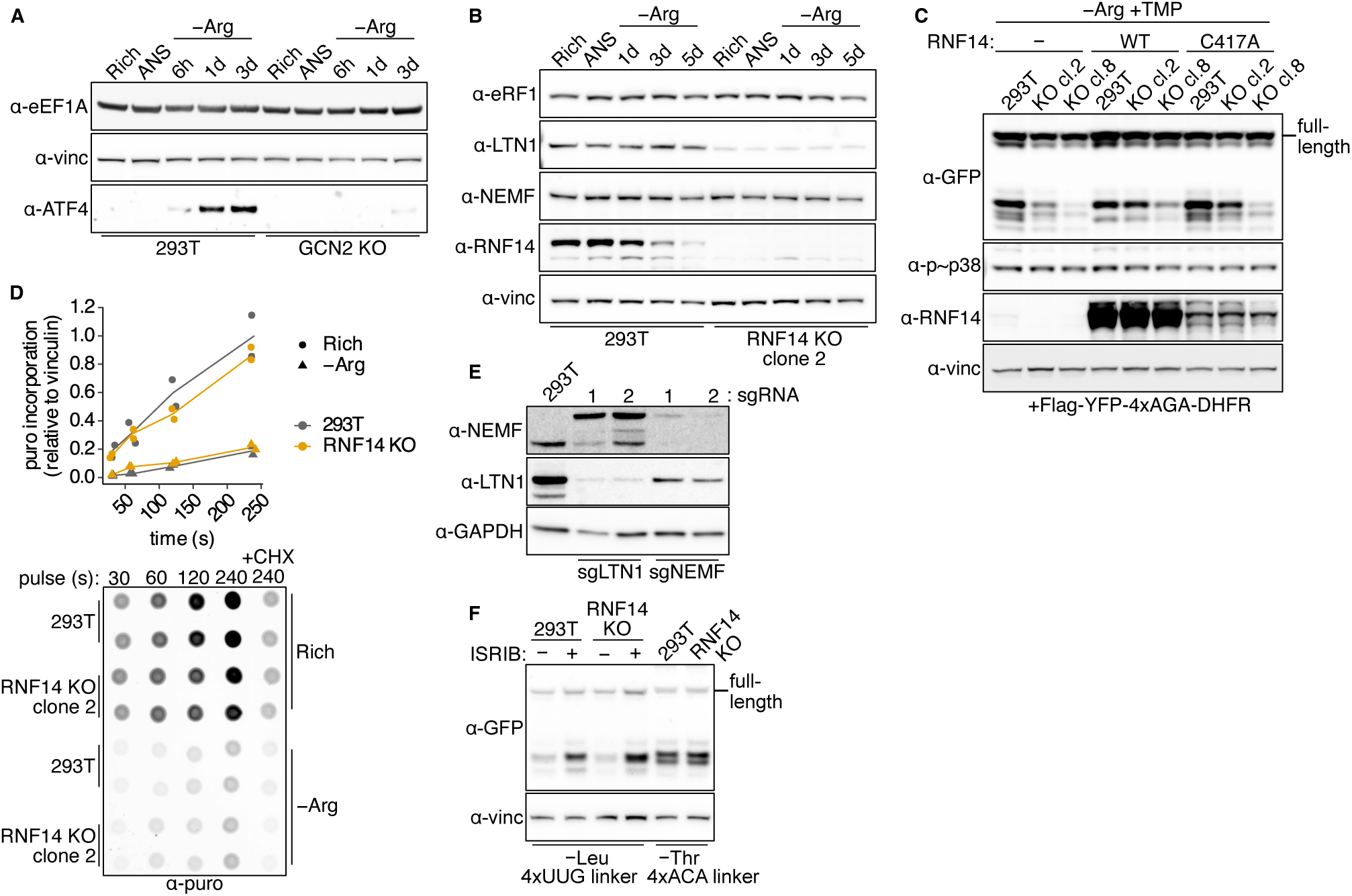
The E3 ubiquitin ligase RNF14 promotes translation disruption at arginine-limited ribosomes. (A-B) Western blots to assess eEF1A (A) or eRF1, LTN1, and NEMF (B) levels with or without GCN2 KO (A) or RNF14 KO (B) over time after arginine limitation or 15 minutes treatment with 0.1 μg/mL anisomycin (ANS) in 293T cells as indicated. (C) Western blot to assess translation disruption product accumulation and p38 phosphorylation in wild-type and RNF14 KO 293T cells expressing nothing, V5-tagged WT, or V5-tagged C417A mutant RNF14 and the Flag-YFP^CGG^-4xAGA-DHFR^CGG^ reporter upon arginine limitation for 3 days, with 10 μM trimethoprim (TMP) to stabilize the degron domain and assess full-length reporter production via read-through. (D) 10 μg/mL puromycin (puro) incorporation into nascent polypeptides after pulsing for the indicated time (s; seconds) in Rich conditions or after 3 days of arginine limitation in wild-type and RNF14 KO 293T cells as indicated. Anti-puro dot blot used for quantification is shown below, including 10 μg/mL cycloheximide (CHX) control added for the last 30 minutes of treatment with arginine limited or rich medium. (E) Western blot to assess LTN1 and NEMF levels in 293T cells after targeting by CRISPRi with 2 independent sgRNAs. (F) Western blot to assess translation disruption product accumulation after 5 days of leucine limitation with or without 40 nM ISRIB or 5 days of threonine limitation in 293T cells with or without RNF14 KO (clone 2) and expressing the Flag-YFP^CUA^-4xUUG-DHFR or Flag-YFP^ACG^-4xACA-DHFR reporters, respectively. (A-C, E, F) Full-length reporter product is indicated (α-GFP antibody used to detect YFP, α-vinc = α-vinculin).

## Supplemental Files

**File S1** – Full flow cytometry time courses for codon-specific protein synthesis rate reporters to select YFP codons; HEK293T (293T), HCT116

**File S2** – Full flow cytometry time courses for translation disruption reporter signal; HEK293T (293T)

**File S3** – Full flow cytometry time courses for translation disruption reporter signal; HCT116, MS1579, MDA-MB-231

**File S4** – Full western blots for translation disruption reporters; HEK293T (293T)

**File S5** – Full western blots for translation disruption reporters; HCT116, MS1579, MDA-MB-231

