## Supplemental Files for "Ribosome-associated quality control of aberrant protein production during amino acid limitation"

**File S1: Full flow cytometry time courses for codon-specific protein synthesis rate reporters to select YFP codons; HEK293T (293T), HCT116**

GCN2 KO = 293T GCN2 KO c2 or HCT116 GCN2 KO c6.
PYCR TKO = triple KO for PYCR isoforms 1, 2, and 3 in HEK293T cells.
TMP = trimethoprim, 10 uM
Fluorescence is plotted relative to baseline measurement in unstarved (Rich) condition for each reporter, dashed line indicates 1. Error bars (where indicated) are standard error of the mean from 2-3 biological replicate measurements.

**Valine: (YFP "safe" codon chosen = GTG), translation defect identified at GTT in GCN2 KO.**

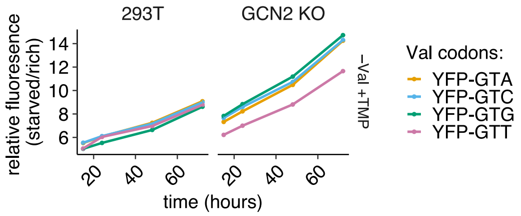


**Proline: (YFP "safe" codon chosen = CCG), translation defect identified at CCA/CCC in GCN2/PYCR TKO. *CCT not tested.**

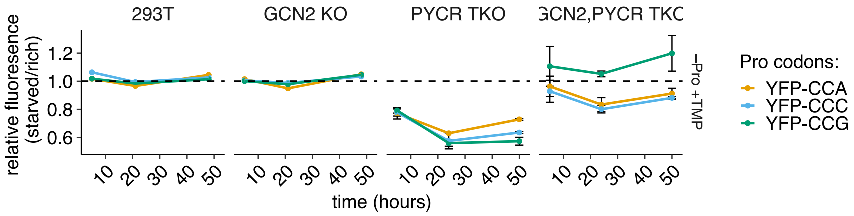


**Alanine: (YFP "safe" codon chosen = GCT). No clear translation defect identified.**

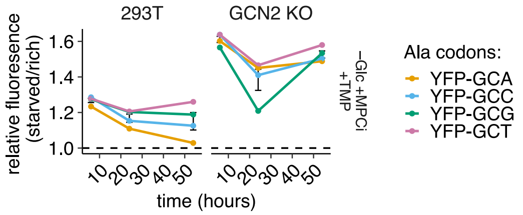


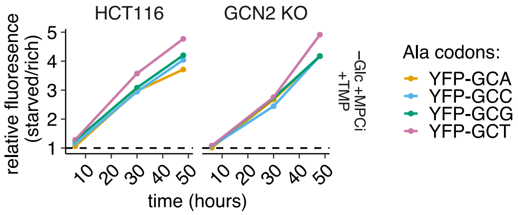


**Glycine: (YFP "safe" codon chosen = GGC). No clear translation defect identified.**

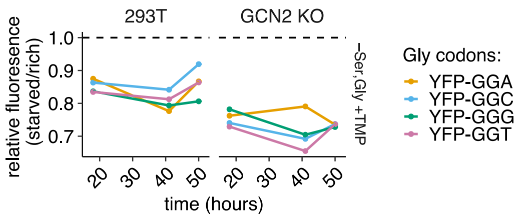


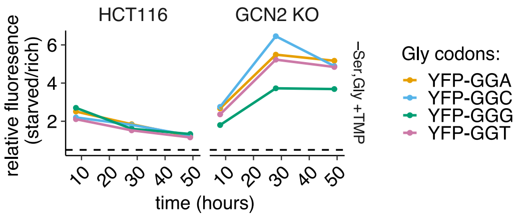


**Threonine: (YFP "safe" codon chosen = ACC), translation defect identified at ACA/ACG in GCN2 KO.**

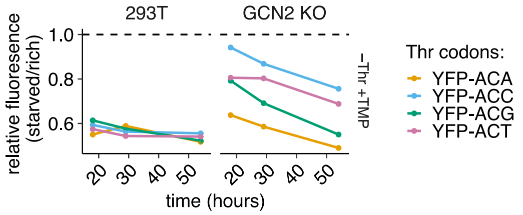


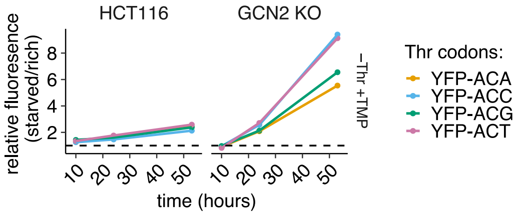


**Arginine (YFP "safe" codon chosen = CGG or CGA; data also shown in Fig S3A), translation defect identified at AGA. GCN2 KO not tested.**

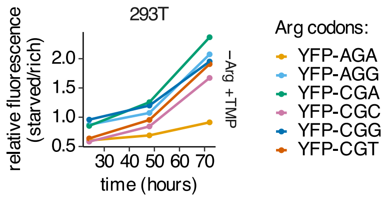


**Isoleucine (YFP "safe" codon chosen = ATA), translation defect identified at ATC/ATT in GCN2 KO.**

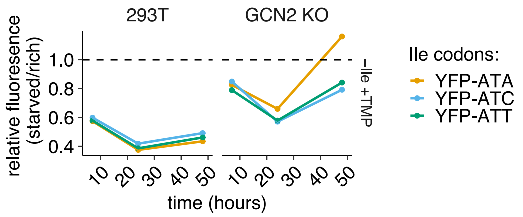


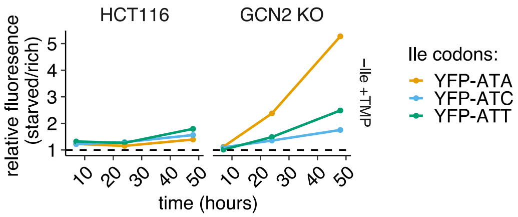

**File S2: Full flow cytometry time courses for translation disruption reporter signal; HEK293T (293T)**

GCN2 KO = 293T GCN2 KO c2. −Glc +MPCi = minus glucose + MPC inhibitor (UK-5099).
Fluorescence is plotted relative to baseline measurement in unstarved (Rich) condition for each reporter, dashed line indicates 1.

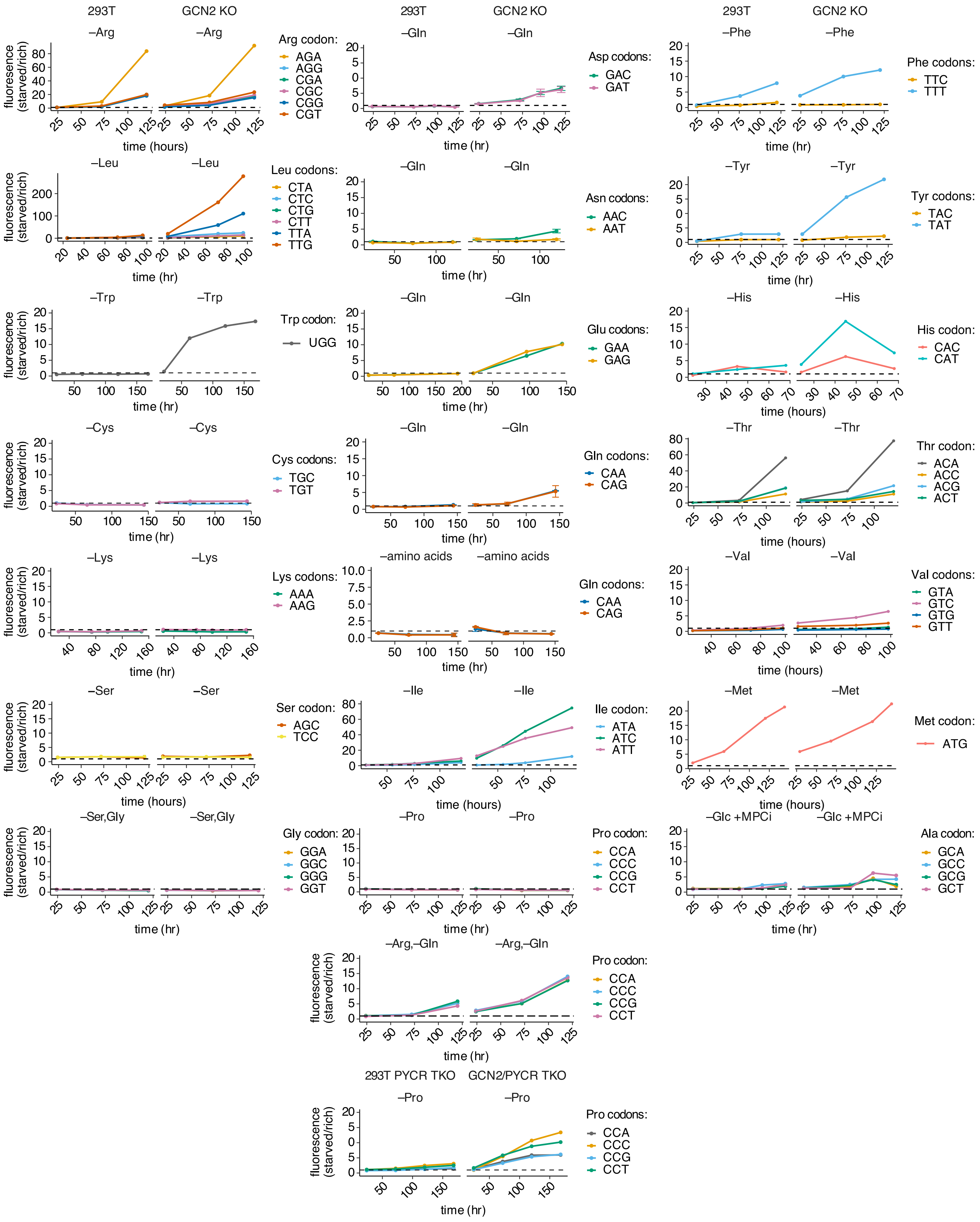

**File S3: Full flow cytometry time courses for translation disruption reporter signal; HCT116, MS1579, MDA-MB-231**

GCN2 KO = HCT116 GCN2 KO c6.
–Glc + MPCi = minus glucose +MPC inhibitor (UK-5099).
Rot = +Rotenone.
Fluorescence is plotted relative to baseline measurement in unstarved (Rich) condition for each reporter, dashed line indicates 1.

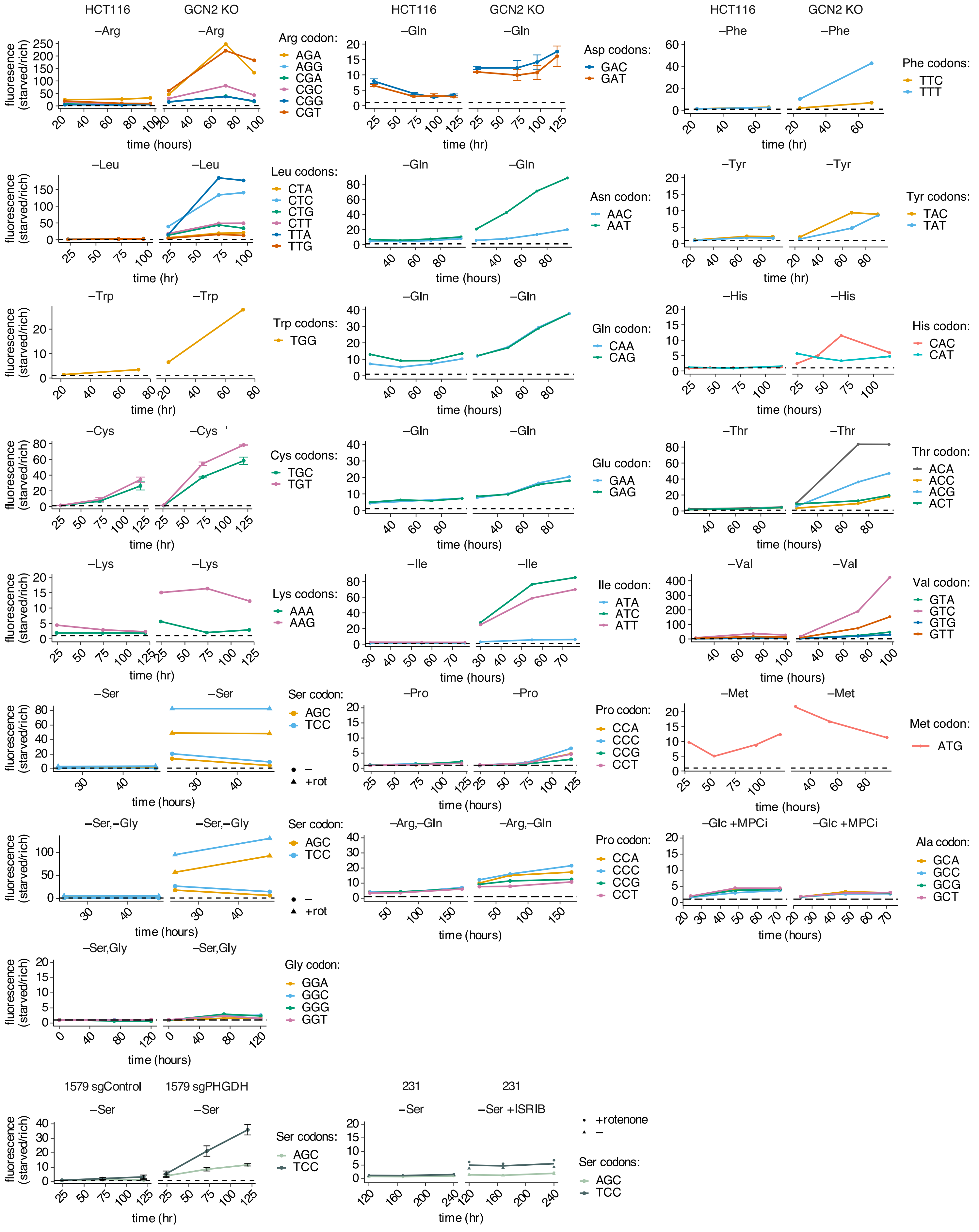

**File S4: Full western blots for translation disruption reporters; HEK293T (293T)**

GCN2 KO = GCN2 KO c2. Rot = +Rotenone.
$Y^{NNN}$;$L^{NNN}$ = YFP encoded with NNN codon variant, Linker to DHFR is 4 NNN (with interspersed Gly codons).
YFP-DHFR = full-length reporter (including N-terminal FLAG)
α-YFP : anti-GFP antibody used to detect YFP. α-vinc : anti-vinculin loading control

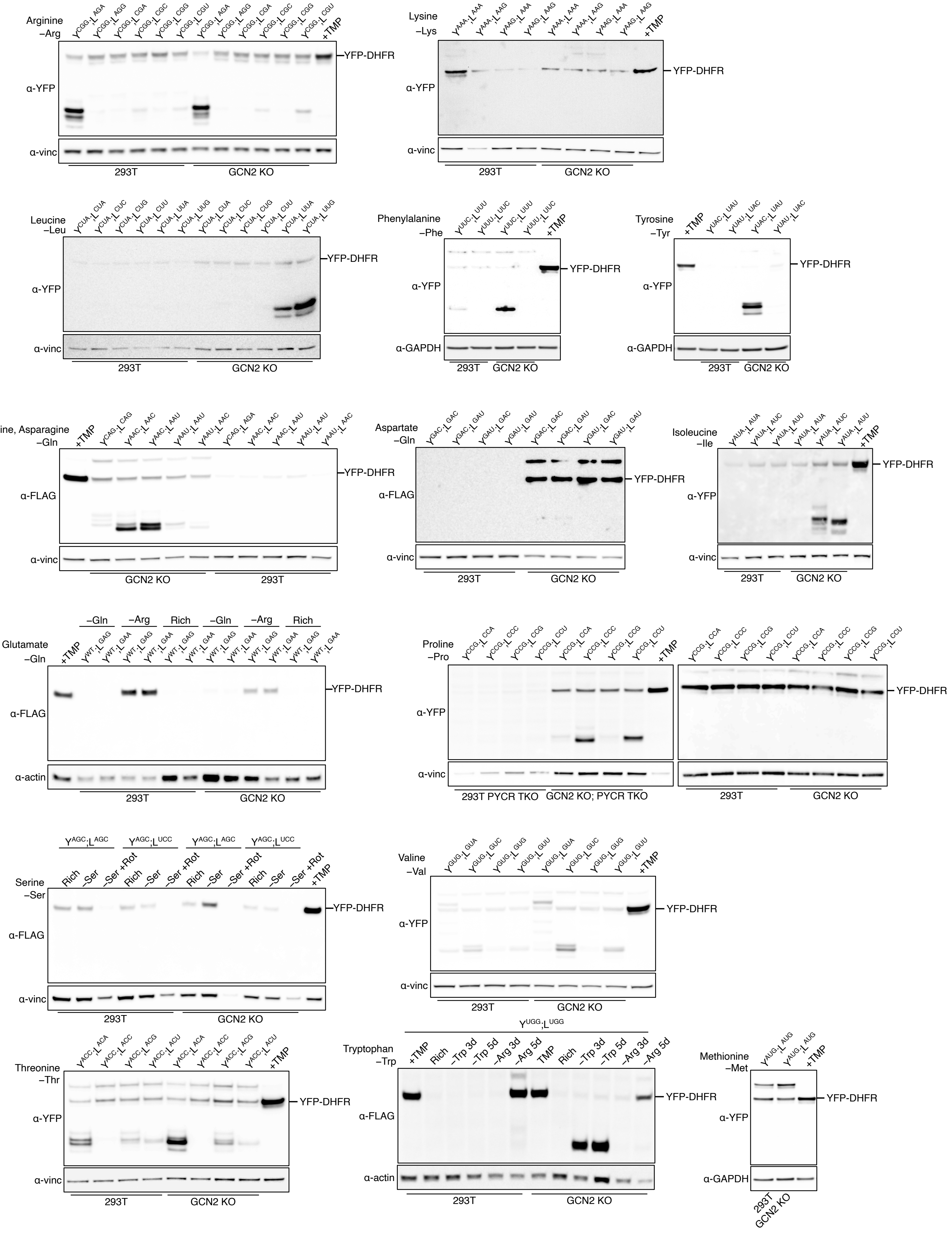

**File S5: Full western blots for translation disruption reporters; HCT116, MS-1579, and MDA-MB-231**

GCN2 KO = HCT116 GCN2 KO c6. Rot = +Rotenone.
$Y^{NNN};L^{NNN}$ = YFP encoded with NNN codon variant, Linker to DHFR is 4 NNN (with interspersed Gly codons).
YFP-DHFR = full-length reporter (including N-terminal FLAG)
α-YFP : anti-GFP antibody used to detect YFP. α-vinc : anti-vinculin loading control

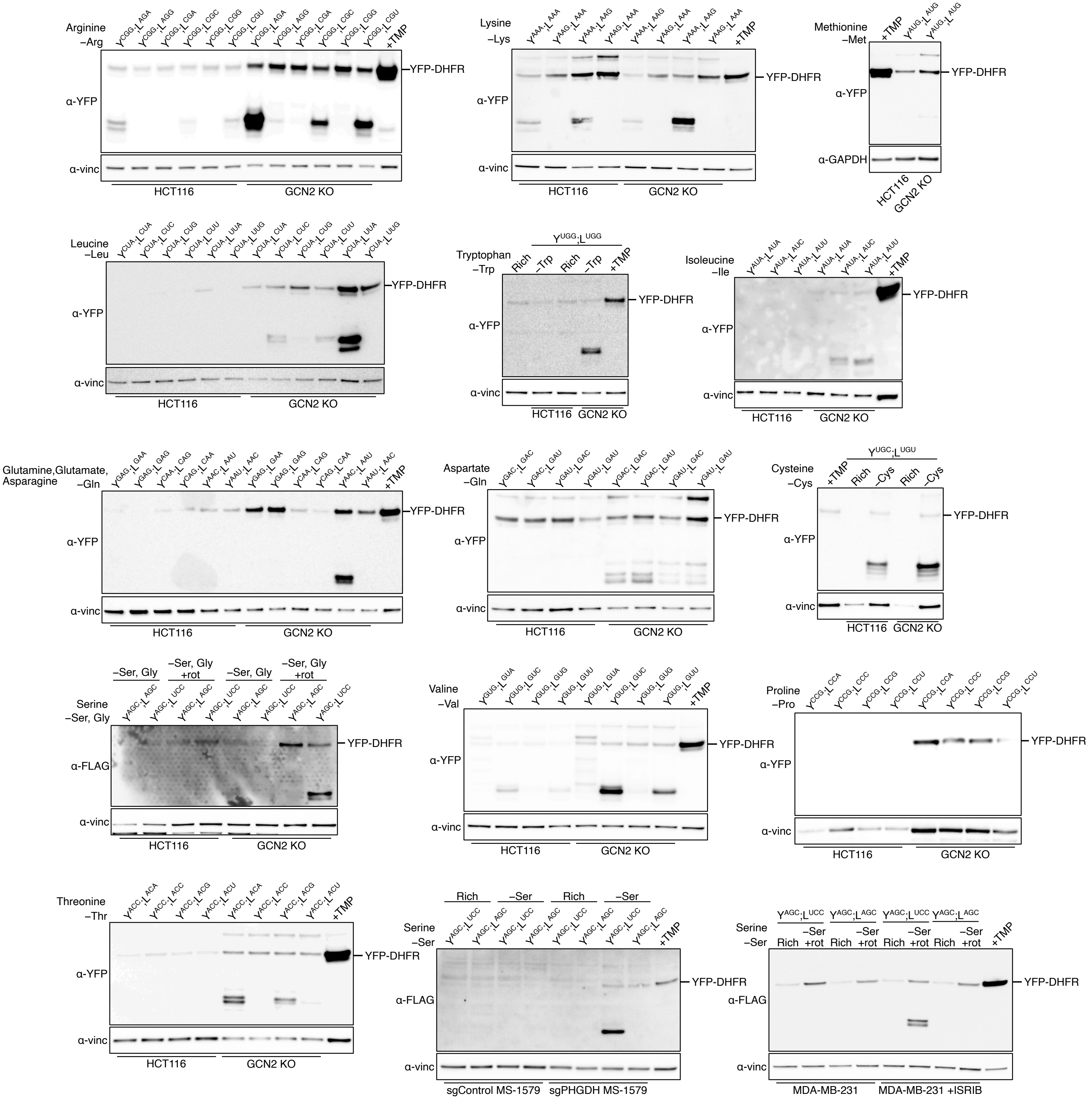
